# Enhancer features that drive formation of transcriptional condensates

**DOI:** 10.1101/495606

**Authors:** Krishna Shrinivas, Benjamin R. Sabari, Eliot L. Coffey, Isaac A. Klein, Ann Boija, Alicia V. Zamudio, Jurian Schuijers, Nancy M. Hannett, Phillip A. Sharp, Richard A. Young, Arup K. Chakraborty

**Affiliations:** Department of Chemical Engineering, Massachusetts Institute of Technology, Cambridge MA 02139.; Institute of Medical Engineering & Science, Massachusetts Institute of Technology, Cambridge MA 02139.; Whitehead Institute for Biomedical Research, 455 Main Street, Cambridge, MA 02142.; Department of Biology, Massachusetts Institute of Technology, Cambridge MA 02139.; Department of Medical Oncology, Dana-Farber Cancer Institute, Harvard Medical School, Boston, MA 02215.; Koch Institute for Integrative Cancer Research, Massachusetts Institute of Technology, Cambridge MA 02139.; Department of Physics, Massachusetts Institute of Technology, Cambridge MA 02139.; Ragon Institute of Massachusetts General Hospital, Massachusetts Institute of Technology, and Harvard University, Cambridge MA 02139.; Department of Chemistry, Massachusetts Institute of Technology, Cambridge MA 02139.

**Author notes:** Correspondence to (P.A.S), (R.A.Y), (A.K.C). Co-first authors (Krishna Shrinivas, Benjamin R. Sabari).

## Abstract

Enhancers, DNA elements that regulate gene expression, contain transcription factor (TF) binding sites. TFs bind short sequence motifs that are present throughout the genome at much higher frequency than active enhancers, and so the features that define active enhancers are not well understood. We show that DNA elements with TF binding site valency, density, and binding affinity above sharply defined thresholds can recruit TFs and coactivators in condensates by the cooperative process of phase separation. We demonstrate that weak cooperative interactions between IDRs of TFs and coactivators in combination with specific TF-DNA interactions are required for forming such transcriptional condensates. IDR-IDR interactions are relatively non-specific with the same molecular interactions shared by many TFs and coactivators, and phase separation is a universal cooperative mechanism. Therefore, whether a genomic locus is an enhancer that can assemble a transcriptional condensate is determined predominantly by its cognate TFs’ binding site valency and density.

## Introduction

The precise regulation of gene transcription during development and in response to signals is established by the action of enhancer elements, which act as platforms for the recruitment of the gene control machinery at specific genomic loci (Levo and Segal, 2014; Long et al., 2016; Maniatis et al., 1998; Ptashne and Gann, 1997; Shlyueva et al., 2014; Spitz and Furlong, 2012). Imprecision in this process can cause disease, including cancer (Herz et al., 2014; Lee and Young, 2013). Enhancer sequences contain short DNA motifs recognized by DNA-binding transcription factors (TFs), which recruit various coactivators that act together to engage RNA Polymerase II (RNAPII) resulting in transcriptional activity (Ptashne and Gann, 1997; Stampfel et al., 2015). Eukaryotic TFs typically recognize short DNA motifs of the order of 6-12 base pairs (Weirauch et al., 2014). There are many such similar affinity motifs in the genome (Lambert et al., 2018; Wunderlich and Mirny, 2009). As a result, active enhancer regions represent only a small fraction of putative binding sites for any given TF (Levo and Segal, 2014; Slattery et al., 2014; Spitz and Furlong, 2012; Wunderlich and Mirny, 2009), with tens of thousands of neutral or non-functional binding events. Determining whether a DNA motif participates in formation of an active enhancer element is thought to require defining a specific set of molecules and the mechanisms by which they act cooperatively to assemble the transcriptional machinery. Because this choice is made from a large set of possibilities, predicting enhancer elements is a significant challenge that has been referred to as the “futility theorem” (Wasserman and Sandelin, 2004).

The presence of clusters of TF binding sites at a genomic locus has been found to be predictive of enhancer elements (Berman et al., 2002; Markstein et al., 2002; Rajewsky et al., 2002). Clusters of TF binding sites can also occur without producing enhancer activity, and enhancer function can be realized upon small insertions (Mansour et al., 2014). The mechanisms by which TF binding site clusters enable the recruitment and stabilization of the appropriate transcriptional machinery at such loci are not well understood. Previous studies into the rules that govern enhancer formation have been focused on cooperativity between TFs, mediated through direct protein-protein interactions or indirectly through changes in chromatin accessibility, nucleosome occupancy and local changes in DNA shape upon binding (Jolma et al., 2015; Lambert et al., 2018; Levo and Segal, 2014; Long et al., 2016; Maniatis et al., 1998; Morgunova and Taipale, 2017; Spitz and Furlong, 2012).

Recent studies suggest that the cooperative process of phase separation can also contribute to assembling dynamic clusters or condensates of TFs, coactivators, and RNA polymerase II at active enhancer elements, especially super-enhancers (Boija et al., 2018; Cho et al., 2018; Chong et al., 2018; Fukaya et al., 2016; Hnisz et al., 2017; Sabari et al., 2018; Tsai et al., 2017). Some of these transcriptional condensates contain hundreds of coactivator and RNA polymerase II molecules (Cho et al., 2018). Many recent studies have shown that multivalent weak interactions between intrinsically disordered regions (IDRs) of proteins promote phase separation, allowing for the compartmentalization and concentration of many biochemical reactions within the cell (reviewed in (Banani et al., 2017; Brangwynne et al., 2015; Shin and Brangwynne, 2017)). Consistent with this picture, TFs and coactivators contain IDRs that promote phase separation and gene activation at enhancers (Boija et al., 2018; Sabari et al., 2018).

If transcriptional condensate formation contributes to assembling certain active enhancers, investigating how features encoded in the DNA element regulate this process should shed light on the cooperative mechanisms that enable the recruitment and stabilization of the transcriptional machinery. The characteristics of DNA sequences that can mediate specific TF-DNA interactions and corresponding weak IDR interactions to form transcriptional condensates at particular genomic loci are unknown.

Using a combination of computational modeling and *in vitro* reconstitutions, we first demonstrate that DNA elements with specific types of TF binding site valency, density, and specificity drive condensation of TFs and coactivators. We study how modulating the affinities, number, or density of TF-DNA interactions and strength of IDR-IDR interactions impacts condensate formation. Because of the cooperative nature of phase separation, condensates form above sharply defined values of these quantities. We then show that the DNA sequence features that promote condensation *in vitro* also promote enhancer activity in cell-based reporter assays. Genome-wide bioinformatic analyses show that these features also characterize known enhancer regions. These results suggest that specific features encoded in DNA elements drive condensate formation, and this may contribute to localization of the transcriptional machinery at enhancers and subsequent enhancer activity. Importantly, we show that a combination of weak multivalent IDR-mediated interactions and strong (specific) TF-DNA interactions are necessary for localized condensation.

The requirement of specific TF-DNA interactions ensures that a transcriptional condensate forms at a locus containing cognate binding sites for the appropriate TF. Although a molecular grammar may describe weak interactions between IDRs, they are relatively non-specific, and the same IDRs bound to different TFs and coactivators are involved in regulating different genes. Furthermore, phase separation is a universal cooperative mechanism for assembling the transcriptional machinery. Consequently, TF binding site valency and density play a dominant role in defining active enhancers formed by a particular TF. Specifically, dense clusters of TF binding sites with a valency exceeding a threshold value are likely to assemble a transcriptional condensate to enable enhancer function. We also report that transcriptional condensate formation can contribute to long-range genomic interactions and organization, potentially promoting enhancer-promoter communication and compartmentalization of actively transcribed regions.

Our study demonstrates that specific features of DNA sequence promote transcriptional condensate formation and enhancer activity, and provides a mechanistic framework to better understand condensate formation at specific genomic loci. Our results also suggest that a general cooperative mechanism contributes to active enhancer formation.

## Results

### Development of a computational model

To explore how the complex interactions between regulatory DNA elements, TFs, and coactivators impact formation of transcription condensates, we first developed a simplified computational model (Figure 1A). The goal of our computational studies was not quantitative recapitulation of experimental data, but rather aimed at obtaining qualitative mechanistic insights that could then be tested by focused experiments. Since enhancers are typically short regions of DNA that are bound by multiple TFs (Levo and Segal, 2014; Spitz and Furlong, 2012), we modeled regulatory DNA elements as a polymer with varying numbers of TF binding sites. Each TF binding site mimics a short (6-12bp) DNA sequence. Specific recognition of DNA motifs by TFs (Weirauch et al., 2014) is mediated by typical TF-DNA binding strengths corresponding to nanomolar dissociation equilibrium constants (Jung et al., 2018), which is the range of TF-DNA interaction energies that we have studied in our simulations (Methods). TFs and coactivators contain large IDRs that interact weakly with each other (Boija et al., 2018; Sabari et al., 2018). Thus, we modeled IDRs of TFs and coactivators as flexible chains attached to their respective structured domains. The IDRs interact with each other via multiple low-affinity interactions. The range of IDR-IDR interaction energies that we have studied in our simulations (Methods) corresponds to those that have been determined by *in vitro* studies of such systems (Brady et al., 2017; Nott et al., 2015; Wei et al., 2017).

**Figure 1:**
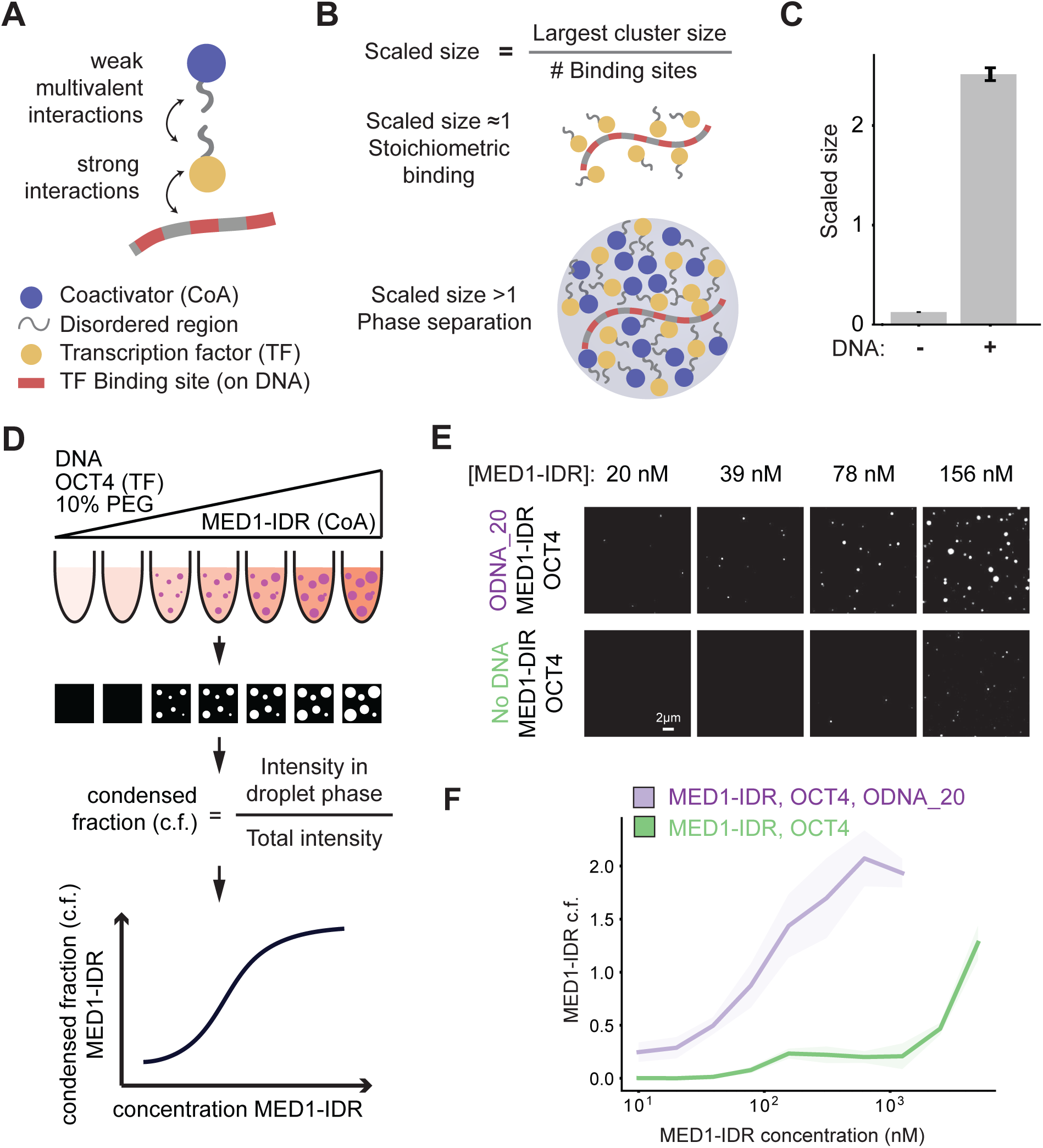
Interactions between TFs and multivalent DNA drives phase separation of TFs and coactivators at low concentrations. A. Schematic depiction of the stochastic computational model and key interactions between molecules. The model consists of a DNA polymer with variable number of TF binding sites, TFs, and coactivators. TFs bind TF binding sites with strong monovalent interactions, and TFs and coactivators interact via weak multivalent interactions between their flexible chains, which mimic the disordered regions of these proteins. B. Scaled size is calculated from simulation trajectories, defined as the size of the largest cluster normalized by the number of DNA binding sites. This value is used as a proxy to differentiate stoichiometric binding of TFs to DNA (scaled size ≈ 1, top illustration) from phase-separated super-stoichiometric assemblies (scaled size >1, bottom illustration). For all reported simulation results, reported quantities are averaged over 10 replicate trajectories. C. Simulations predict that multivalent DNA-TF interactions result in phase separation of TF and coactivator at dilute concentrations, as shown by scaled size >1 upon addition of DNA. D. Schematic depiction of experimental workflow and image analysis for *in vitro* droplet assay. DNA, OCT4, and varying concentrations of MED1-IDR are incubated together in the presence of 10% PEG-8000 as a molecular crowder (illustrated with test tubes, see methods for detail). Fluorescence microscopy of these mixtures is used to detect droplet formation (illustrated by black square with or without white droplets). Multiple images per condition are then analyzed to calculate condensed fraction (c.f.) as intensity of fluorescence signal within droplets divided by total intensity in the image. Condensed fraction over MED1-IDR concentration is graphed (illustrated by a cartoon graph). For each experimental condition (e.g. varying type or presence of DNA or OCT4) c.f. curves are compared to see how the experimental condition impacts coactivator phase separation. E. Representative images of MED1-IDR droplets in the presence of OCT4 and ODNA_20 (top row) or with only OCT4 (bottom row) at indicated MED1-IDR concentrations. See Table S2 for sequence of DNAs used in droplet assays. F. Condensed fraction of MED1-IDR (in units of percentage) with DNA (purple) or without DNA (green) across a range of MED1-IDR concentrations. Higher condensed fraction implies higher fraction of total signal in droplet phase. Solid lines represent mean and shaded background represents boundaries of mean±std. See methods for details on calculation of condensed fraction.

We simulated this model using standard Langevin molecular dynamics methods to calculate spatiotemporal trajectories of the participating species (see methods, (Anderson et al., 2008)). To distinguish stoichiometrically bound complexes from a phase-separated condensate, we computed the size of the largest molecular cluster scaled by the number of TF binding sites present on DNA. Values of this scaled size greater than 1 represent phase-separated super-stoichiometric assemblies (i.e. condensates), while values close to 1 correspond to stoichiometrically bound TFs (Figure 1B). Using the scaled size as a measure of transcriptional condensate formation, we studied how particular motif compositions on DNA, as well as TF-DNA interactions and interactions between TF and coactivator IDRs regulate transcriptional condensate formation at DNA loci.

### Interactions between TFs and multivalent DNA drive formation of condensates of TFs and coactivators

TFs and coactivators form condensates *in vitro* at supra-physiological concentrations (Boija et al., 2018; Lu et al., 2018; Sabari et al., 2018). Our simulation results (Figure 1C) predict that a dilute solution of TFs and coactivators that does not phase separate by itself forms condensates (scaled size greater than 1) upon adding multivalent DNA (DNA with 30 TF binding sites). To test this prediction, we developed an *in vitro* phase separation droplet assay containing the three components present in our simulations: TF, coactivator, and DNA (Figure 1D). For TF and coactivator, we employed purified OCT4, a master transcription factor in murine embryonic stem cells (mESCs), and MED1-IDR, the intrinsically disordered region of the largest subunit of the Mediator coactivator complex. We have previously shown that these proteins phase separate together *in vitro* and *in vivo* (Boija et al., 2018; Sabari et al., 2018). For DNA, we used various synthetic DNA sequences containing varying numbers of OCT4 binding sites (see methods and Table S2). Each of the three components was fluorescently labeled either by fluorescent protein fusion, mEGFP-OCT4 and mCherry-MED1-IDR, or a fluorescent dye, Cy5-DNA.

Formation of phase-separated droplets was monitored over a range of MED1-IDR concentrations by fluorescence microscopy with a fixed concentration of OCT4 in the presence or absence of multivalent DNA (DNA with 20 OCT4 binding sites, 8bp motif with 8bp spacers, ODNA_20, see methods and Table S2). The fluorescence microscopy results were quantified by calculating the condensed fraction (Figure 1D, also see methods). Consistent with model predictions, addition of DNA promoted phase separation at low MED1-IDR concentrations (Figure 1E). Graphing the condensed fraction over a range of MED1-IDR concentrations further validated that the addition of ODNA_20 promoted phase separation at low MED1-IDR concentrations (Figure 1F). These results demonstrate that multivalent DNA promotes the phase separation of TFs and coactivators at low protein concentrations, comparable to concentrations observed *in vivo* (Figure S1A).

To further study how DNA influences condensate stability, we performed simulations where TF and coactivator condensates were allowed to form in the presence of DNA, followed by a simulated disruption of TF-DNA interactions. At dilute protein concentrations, disrupting TF-DNA interactions resulted in dissolution of condensates (Figure 2A orange line, Movie S1), demonstrating that, under these conditions, DNA is required for both formation and stability of condensates. While addition of DNA at high protein concentrations increased the rate of condensate assembly by acting as a seed (Figure S2; Movie S2), disruption of TF-DNA interactions at these high concentrations did not lead to condensate dissolution (Figure 2A; grey line). We observed a drop in scaled size upon TF-DNA interaction disruption in this case, but this was due to the condensate being broken into smaller droplets as the DNA was ejected from the condensate (as depicted in Figure 2A; grey box, Movie S2). These results predict that, at dilute protein concentrations, specific TF-DNA interactions are required for both formation and stability of condensates at particular genomic loci with DNA acting as a scaffold for phase separation.

**Figure 2:**
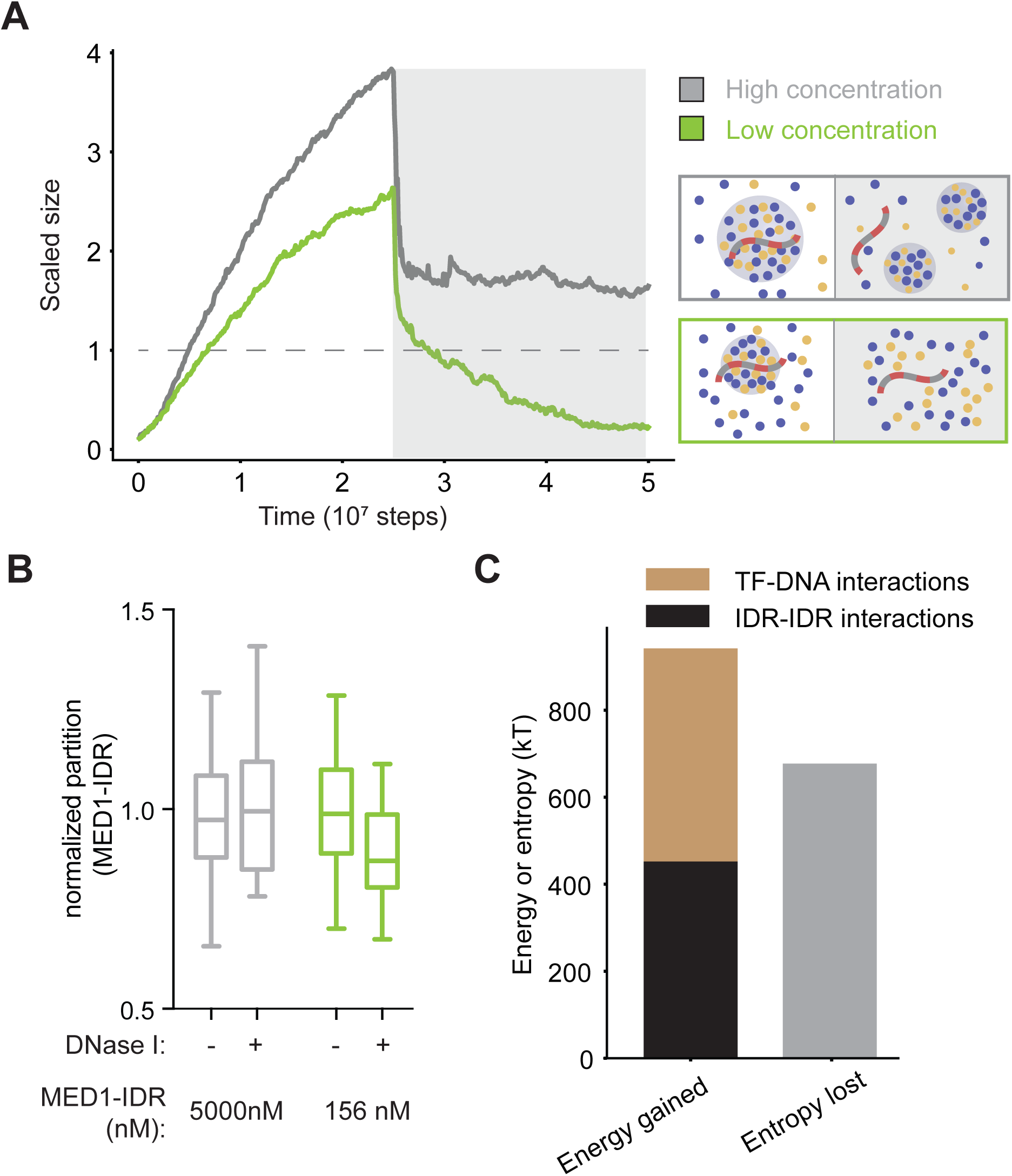
Transcriptional condensate stability is governed by a combination of TF-DNA and IDR-IDR interactions between TFs and coactivators. A. Simulation results for dynamics of condensate assembly/disassembly at two different protein concentrations is represented by average scaled size on the ordinate, and time (in simulation steps after initialization) on the abscissa. TF-DNA interactions are disrupted after stable condensate assembly (shown by a dark gray background). Schematic depiction of phase behavior is shown enclosed in boxes whose colors match the respective lines. See Movies S1 and S1. B. Box-plot depiction of experimentally determined MED1-IDR partition ratio (see methods) between condensate and background, at high (5000nM, gray) and low concentrations (156nM, green) of MED1-IDR in the presence of OCT4 and ODNA_20, in the absence (-) or in the presence (+) DNase I. The partition ratio is normalized to the (-) condition, and lower partition ratios imply lesser enrichment of MED1 in the droplet phase. C. Energetic attractions, arising from a combination of TF-DNA (brown) and IDR (black) interactions, compensate for entropy loss (grey) of forming a condensate.

To mimic disruption of TF-DNA interactions *in vitro*, we added DNase I to droplets formed at high or low concentrations of MED1-IDR in presence of OCT4 and ODNA_20 (see Figure 2 caption and methods). As expected, DNA was degraded in both conditions (Figure S1B), as measured by a decrease in the enrichment of DNA in droplets (see methods). Consistent with our model predictions, droplets formed at the lower concentrations were more sensitive to the degradation of DNA than were those formed at higher concentrations (Figure 2B). While enzymatic degradation of DNA did not completely dissolve droplets in our *in vitro* experiments, MED1-IDR enrichment within droplets was affected only at the lower protein concentration (Figure 2B). Together, the *in silico* and *in vitro* results indicate that DNA can nucleate and scaffold phase-separated condensates of TFs and coactivators.

### Physical mechanisms that underlie localized formation of transcriptional condensates

To understand the mechanisms driving DNA-mediated condensate formation, we cast our results in terms of the competing thermodynamic forces that govern phase separation. For computational efficiency, further characterization of our model was carried out with a simplified implicit-IDR model (Figure S3A), which recapitulated all features (Figure S3B) of the explicit-IDR model. Typically, condensate formation results in entropy loss because the molecules in the droplet are more confined than if they were in free solution. A condensate is stable only if this entropy loss is compensated by the enhanced attractive interaction energies between molecules confined in the condensate. We computed the potential energy by summing up all pairwise molecular interactions. Entropy loss due to confinement was calculated by adding a factor of 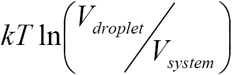 for each molecule in the condensate. Our simulations show that a sum of specific TF-DNA interactions and weak IDR interactions between TFs and coactivators are necessary to compensate for the entropy loss of forming condensates at low concentrations (Figure 2C). IDR interactions alone are insufficient to compensate for the entropy loss of condensate formation, thus disruption of TF-DNA interactions results in condensate dissolution (Figures 2 and S3B-S3C, dark background). Likewise, TF-DNA interactions alone are insufficient to compensate for the entropy loss of condensate formation and disruption of IDR-IDR interactions results in condensate dissolution (see next section). The same features are observed in explicit-IDR simulations (Figures 2A; orange line, and S3D), though our simplified calculation of the entropy loss in this case (see above) is an underestimate as contributions from the change in configurational entropy of IDR chains is not accounted. These results provide a mechanistic framework to understand how the combination of TF-DNA interactions and weak IDR interactions determine assembly and stability of transcription condensates at low concentrations.

### Specific TF-DNA interactions and weak multivalent IDR interactions regulate formation of transcriptional condensates

Given that TF-DNA interactions are necessary for condensate formation, we next investigated the effect of modulating TF-DNA affinity at dilute protein concentrations. Simulations predict that condensates form above a sharply defined affinity threshold (Figure 3A), and that high affinity TF-DNA interactions result in condensate formation at low coactivator concentration thresholds (Figure 3B). Using the *in vitro* droplet assay, we probed the effect of TF-DNA interactions by comparing phase separation of MED1-IDR over a range of concentrations, with fixed concentrations of both OCT4 and either ODNA_20 or a scrambled ODNA_20 which does not contain any consensus binding sites for OCT4 (ODNA_20sc, Table S2). High-affinity OCT4-ODNA_20 interactions promoted phase separation at lower MED1-IDR concentration when compared to OCT4-ODNA_20sc (Figure 3C), further corroborated by quantifying the condensed fraction of MED1-IDR over the concentrations tested (Figure 3D). The same lowered concentration threshold was observed when quantifying the condensed fraction of OCT4 or DNA (Figure S4A-B). These results demonstrate that higher TF-DNA affinities promote phase separation above sharply defined thresholds. Therefore, TFs, which exhibit only a 10-100 fold higher affinity for specific DNA binding sites compared to random DNA, can drive transcriptional condensate formation at specific DNA loci.

**Figure 3:**
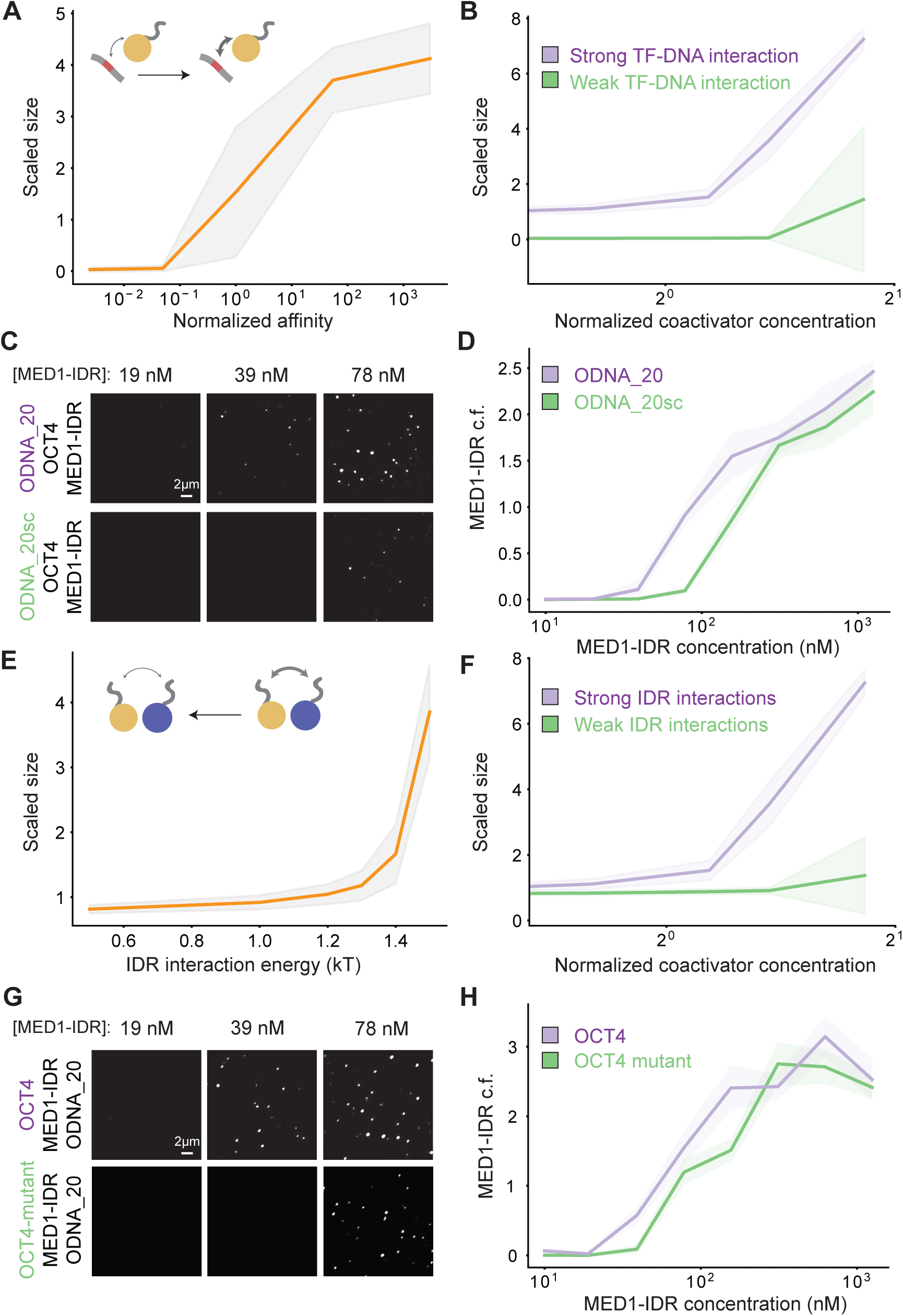
Phase separation is regulated through strong specific TF-DNA interactions and weak multivalent interactions between TF and coactivator IDRs. A. Simulations predict a shift in scaled size from stoichiometric binding (≈1) to phase separation (>1) with increasing normalized affinity (darker arrow in schematic); affinity normalized to threshold affinity of E=12kT. B. Scaled size predictions for high (normalized affinity ≈50, purple) and low (normalized affinity ≈5e-2, green) TF-DNA affinities as a function of coactivator concentration. Coactivator concentrations are normalized to value of *N*_*coA*_ = 150. C. Representative images of MED1-IDR droplets with OCT4 and ODNA_20 (top row) or ODNA_20sc (bottom row) at indicated MED1-IDR concentrations. See Table S2 for sequence of DNAs used in droplet assays. D. Condensed fraction of MED1-IDR (in units of percentage) for ODNA_20 (purple) and ODNA_20sc (green) across a range of MED1-IDR concentrations. Higher condensed fraction implies higher fraction of total signal in droplet phase. E. Simulations predict a shift in scaled size from phase separation (>1) to stoichiometric binding (≈1) upon decreasing IDR interaction (from right to left, lighter arrow in schematic). F. Scaled size predictions for high (IDR = 1.5kT, purple) and low (IDR=1.0 KT, green) IDR interaction as a function of coactivator concentration (normalized as in 3B). G. Representative images of MED1-IDR droplets with ODNA_20 and OCT4 (top row) or OCT4-mutant (bottom row) at indicated MED1-IDR concentrations. H. Condensed fraction of MED1-IDR (in units of percentage) for OCT4 (purple) and OCT4-mutant (green) across a range of MED1-IDR concentrations. In all line plots, solid lines represent mean and shaded background represents boundaries of mean±std. See methods for details on calculation of condensed fraction.

We next investigated the effect of modulating the multivalent IDR interactions, whose effective affinity *in vivo* can be regulated through post-translational modifications (Banani et al., 2017; Shin and Brangwynne, 2017). Reducing the strength of IDR interactions between TFs and coactivators in our simulations predicts that condensates dissolve below a sharply defined interaction threshold (Figure 3E), and strong IDR interactions result in condensate formation at lower coactivator concentration thresholds (Figure 3F). To test this prediction, we monitored MED1-IDR phase separation over a range of MED1-IDR concentrations with fixed concentrations of both ODNA_20 and either OCT4 or a previously characterized OCT4 activation-domain mutant (acidic to alanine mutant) with reduced interaction with MED1-IDR (Boija et al., 2018). Consistent with simulation predictions, the OCT4 mutant was less effective at promoting phase separation at low concentrations, as compared to OCT4 (Figures 3G-H). These results further highlight the importance of the interactions of coactivator IDRs with TF IDRs in the formation of transcriptional condensates.

Our results thus far suggest the following model. Specific TF-DNA interactions localize TFs to particular genomic loci. Transcriptional condensate formation is a cooperative process, which occurs at these loci when interactions between the IDRs of TFs and coactivators also exceed a threshold. While other processes are also involved (e.g. DNA bending, removal and modification of nucleosomes, and interactions with RNA), this cooperative phenomenon of TF and coactivator phase separation contributes to assembling the transcriptional machinery at enhancers that form condensates.

### Specific motif compositions encoded in DNA facilitate localized transcriptional condensate formation

To begin defining the specific DNA sequence features that result in condensate formation, we explored the effects of modulating the valency and density of TF binding sites with the same TF-DNA affinities. We reasoned that the same energetic compensation for entropy loss we observed by increasing TF-DNA affinities (Figures 2C and 3A-D) could be obtained instead through increasing the number of DNA binding sites (i.e., valency). Our simulations predict that, for the same TF-DNA binding energies, condensates form above a sharply defined valency threshold (Figure 4A), and higher valency results in condensate formation at lower coactivator concentrations (Figure 4B). Consistent with this prediction, *in vitro* assays revealed that ODNA_20 promoted phase separation of MED1-IDR and OCT4 at a lower concentration threshold than DNA with fewer binding sites (ODNA_5) (Table S2) (Figures 4C-4D).

**Figure 4:**
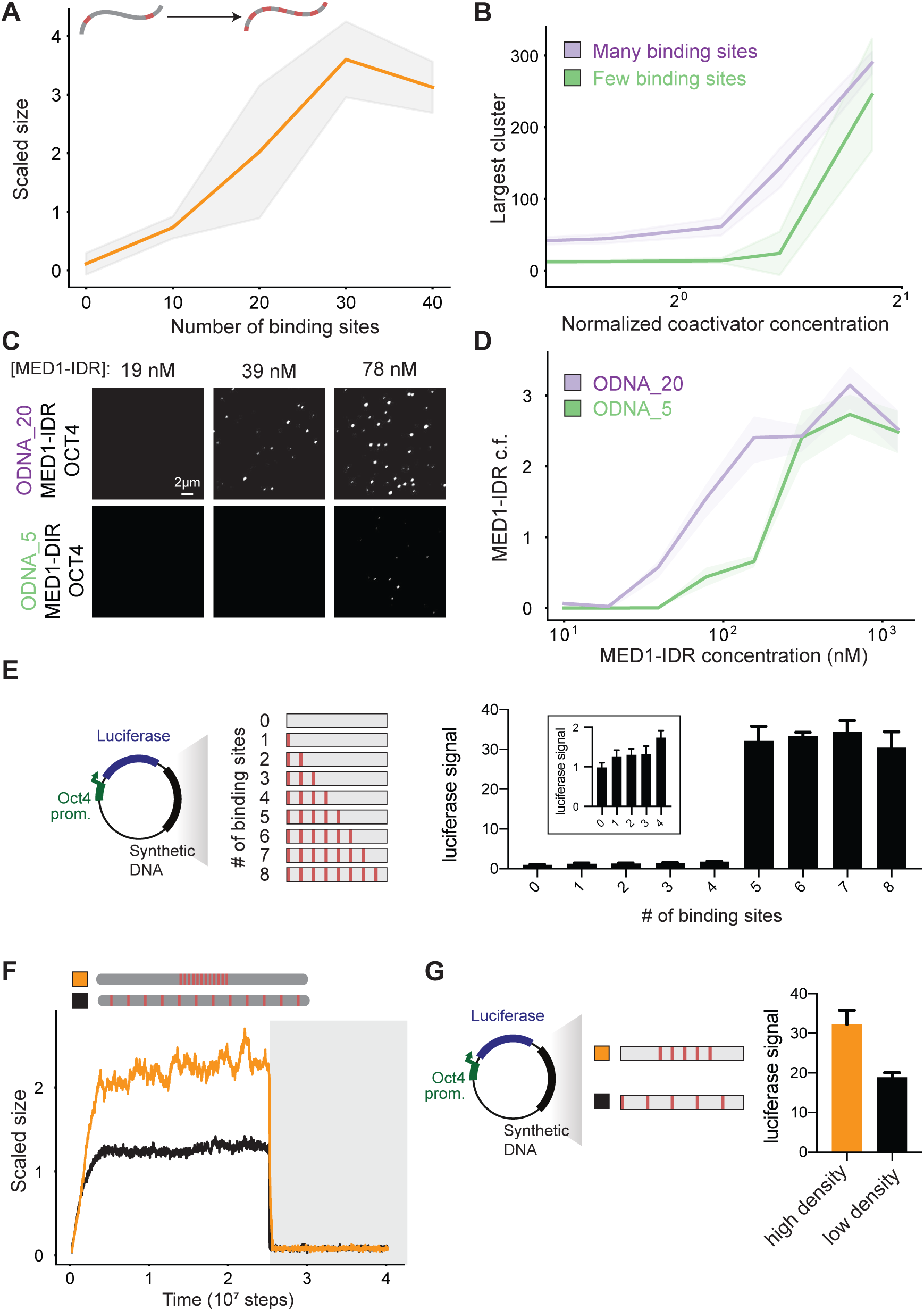
Motif valency and density encoded in DNA drives phase separation. A. Simulations predict a shift in scaled size from stoichiometric binding (≈1) to phase separation (>1) with increasing number of TF binding sites on DNA (schematic depicts increasing number of binding sites). B. Scaled size predictions for many (N =30, purple) and few (N=10, green) binding sites as a function of coactivator concentration (normalized as in 3B). C. Representative images of MED1-IDR droplets with OCT4 and ODNA_20 (top row) or ODNA_5 (bottom row) at indicated MED1-IDR concentrations. See Table S2 for sequence of DNAs used in droplet assays D. Condensed fraction of MED1-IDR (in units of percentage) for ODNA_20 (purple) and ODNA_5 (green) across a range of MED1-IDR concentrations. E. Enhancer activity increases over a sharply defined TF binding site threshold. The left panel shows a schematic depiction of the luciferase reporter construct and the synthetic DNA sequences tested. The right panel shows the luciferase signal from constructs with the indicated number of binding sites transfected into murine embryonic stem cells (see methods). Inset presents data for 0-4 binding sites graphed on a different scale for the ordinate. Luciferase signal is normalized to the construct with 0 motifs. Data is graphed as average of three biological replicates ± std. See Table S3 for sequence of DNAs used in luciferase reporter assays F. Scaled size versus simulation time steps comparing three different distribution of binding site densities (shown in the schematic below). TF-DNA interactions are disrupted after stable condensate assembly (dark gray background). G. DNA sequences with the same number of binding sites, but higher density, shows increase in transcription activity. Left half shows a schematic depiction of the luciferase reporter construct and the synthetic DNA sequences tested. The right half shows the luciferase signal from constructs with indicated binding site density transfected into mouse embryonic stem cells. Data graphed as in E. In all line plots, solid lines represent mean and shaded background represents boundaries of mean±std. See methods for details on calculation of condensed fraction.

To test how motif valency impacts enhancer activity in cells, we cloned synthetic DNA sequences with varying number of OCT4 binding sites into previously characterized luciferase reporter constructs (Whyte et al., 2013) that were subsequently transfected into mESCs(see methods and figure 4E schematic). In these reporter assays, expression of the luciferase gene, read out as luminescence, is a measure of the strength of enhancer activity. Using a series of DNAs with 0 to 8 binding sites (8bp motif with 24bp spacers, see Table S3) we found that enhancer activity increased above a sharply defined valency threshold (Figure 4E), in striking qualitative agreement with simulation predictions for condensate formation (Figure 4A).

To distinguish whether this behavior stemmed from motif valency alone or local motif density, we carried out simulations of DNA chains with a fixed number of binding sites, but different distributions along the chain (Figure 4F). We found that high local density, not total number, of binding sites promoted condensate formation (Figure 4F). *In vitro* experiments with DNA of same binding site number, but different densities, validated this prediction (Figure S4C). To test the effect of binding site density on enhancer activity in cells, we compared the enhancer activity of 5 binding sites with 24bp spacers (high density) to 5 binding sites with 56bp spacers (low density) in luciferase assays in mESCs (Figure 4G). In agreement with the model predictions, reducing density of binding sites led to reduced enhancer activity.

The results in Figure 4 show that dense clusters of a particular TF’s binding sites, with the valency of binding sites exceeding a sharply defined threshold, drive localized formation of transcriptional condensates, and that these same features influence enhancer activity in cells. The condensates form by the universal cooperative mechanism of phase separation which, in turn, requires weak cooperative interactions between the IDRs of TFs and coactivators (Figure 3). IDR-IDR interactions are relatively non-specific, and the same coactivator IDRs can assemble at different enhancers upon cognate TF binding. Therefore, a TF’s binding site valency and density (or binding site clustering) predominantly define its active enhancers since this enables a universal cooperative mechanism to assemble the transcriptional machinery at different genomic loci via similar molecular interactions.

### Transcriptional condensate formation may facilitate long-range interactions and higher-order genome organization

Given that regulatory elements often communicate over long linear distances, we next investigated whether two dense clusters of TF binding sites in DNA separated by a linker could assemble a single condensate. Our simulations show that this is indeed the case (Figure 5A; green line). Hi-C maps computed from the simulation data (see methods) show long-range interactions between the dense clusters of binding sites, which are absent (Figure 5B) at conditions with a low density of TF binding sites distributed uniformly (Figure 5A; black line). These results suggest that condensate formation could explain recent observations of CTCF/cohesin-independent long-range interactions between active regions of the genome (Rowley et al., 2017; Schwarzer et al., 2017). More generally, our results suggest that localized transcriptional condensate formation can facilitate higher-order organization of the 3D-genome and contribute to long-range communication between enhancer-promoter pairs.

**Figure 5:**
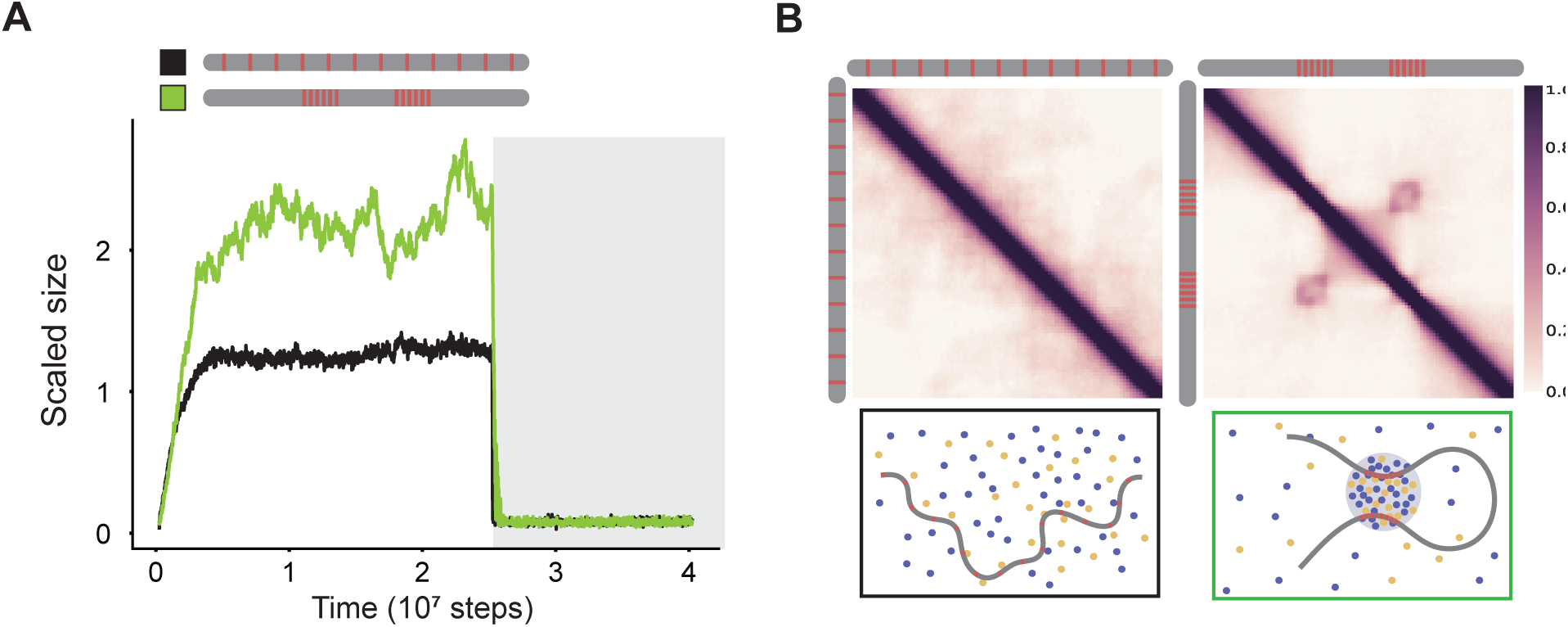
Transcriptional condensate formation facilitates long-range interactions. A. Scaled size versus simulation time steps comparing two different distribution of binding site densities (as shown in the schematic legend). TF-DNA interactions are disrupted after stable condensate assembly (as shown in dark gray background). B. Simulated Hi-C maps (see methods) show long-range interactions (right panel, checkerboard-like patterns) for high local motif density (computed for green line in Figure 5A), and not for low motif density (left panel, computed for black line in Figure 5A). Illustrations depicting the organization of model components are provided for each condition below their respective Hi-C map.

### Mammalian genomes encode specific motif features in enhancers to assemble high densities of transcription apparatus

We next investigated whether enhancer features that our results suggest promote transcriptional condensate formation are present in mammalian genomes. Given that our results show that a linear increase in TF binding site valency can result in an exponential increase in coactivator recruitment by condensate formation (Figure 4), we investigated the relationship between TF binding site valency (i.e. occurrence of TF motifs) and coactivator recruitment in mESCs. We gathered genome-wide distribution of TF motif occurrence for highly expressed mESC master TFs – OCT4, SOX2, KLF4, ESRRB (OSKE). Super-enhancers, genomic regions with unusually high densities of transcriptional molecules (Whyte et al., 2013), where transcriptional condensates have recently been observed (Boija et al., 2018; Cho et al., 2018; Sabari et al., 2018), have higher OSKE motif densities when compared to typical enhancers or random loci (Figures 6A-B, methods). Consistent with our results, we found a highly non-linear (roughly exponential) correlation between OSKE motif density and ChIP-Seq data for MED1, RNA Pol II (Figure 6C), and BRD4 (Figure S5A) across genetic regions including SEs, TEs, and random loci. This correlation was minimal when input control data was analyzed (Figure S5B). These results suggest that enhancer elements that encode specific DNA sequence features can recruit unusually high densities of transcriptional apparatus by transcriptional condensate formation in mammalian genomes, consistent with our results. The same features enable recruitment of varied cofactors – BRD4, MED1, and Pol II, thus suggesting that phase separation contributes to stabilization of transcription machinery at specific genomic loci.

**Figure 6:**
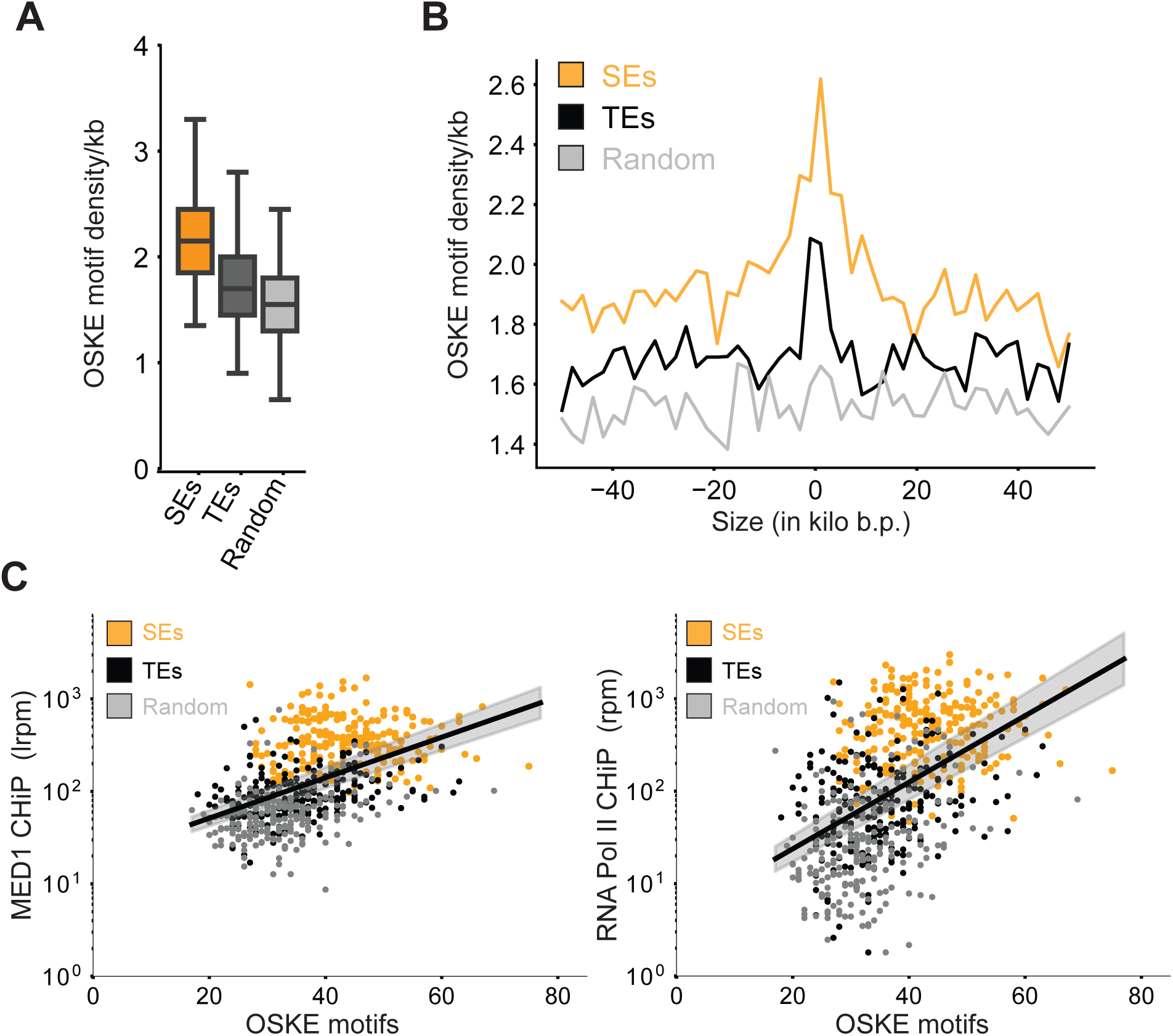
Mammalian genomes leverage high motif density to assemble high density of transcriptional apparatus at key regulatory elements. A. Box-plot depiction of motif density (per kb) of master mESC TFs – OCT4 + SOX2 + KLF4 + ESRRB (OSKE), over 20kb regions centered on super-enhancers (SEs, orange), typical enhancers (TEs, black), and random loci (light gray). B. OSKE motif density over a 100kb window centered at SEs (orange), TEs (black), and random loci (gray). C. MED1 (left) and RNA Pol II (right) ChIP-Seq counts (ordinate, reads-per-million) against total OSKE motifs over 20kb regions centered on SEs (orange), TEs (black), and random loci (gray). The black line is a fit inferred between the logarithmic ChIP signal and the linearly graphed motif count across all regions, and so the fit represents a highly non-linear (exponential) correlation. The grey shaded regions represent 95% confidence intervals in the inferred parameters. The exponential fit describes a sizable fraction of the observed variance i.e. *R*^2^ ≈ 0.25 for both inferred lines.

## Discussion

Enhancers are DNA elements that control gene expression by assembling the transcriptional machinery at specific genomic loci. Active enhancers are considered to form by cooperative interactions between the molecular participants and with the DNA element. Defining a specific combination of molecules and the mechanisms of cooperative interactions between them that enables assembly of the transcriptional machinery at a particular enhancer is challenging since this choice is made from a large number of possibilities. Studies addressing this challenge provide insights into the mechanisms that describe how enhancers form to regulate transcription, and how aberrant regulation leads to disease.

Past studies have largely focused on cooperative mechanisms that include specific protein-protein interactions, DNA-bending mediated protein binding, and modification of nucleosomes (Jolma et al., 2015; Lambert et al., 2018; Levo and Segal, 2014; Long et al., 2016; Maniatis et al., 1998; Morgunova and Taipale, 2017; Spitz and Furlong, 2012). While such mechanisms are undoubtedly at play, our results suggest that another more general mechanism may contribute to assembling the transcriptional machinery at active enhancers.

Recent studies have suggested that phase-separated condensates of molecules involved in transcription form at enhancers (Boija et al., 2018; Cho et al., 2018; Chong et al., 2018; Fukaya et al., 2016; Hnisz et al., 2017; Sabari et al., 2018; Tsai et al., 2017), providing a potential mechanism for concentrating transcriptional machinery at specific loci. Here we investigated how features encoded in a DNA element regulate the formation of transcriptional condensates. Our results shed light on the features of enhancer sequences that can enable assembly of the transcriptional machinery by the general cooperative mechanism of phase separation.

We first demonstrated that interactions between TFs and multivalent DNA elements can form condensates of TFs and coactivators at protein concentrations that are too low for these molecules to phase separate by themselves. This result may explain why condensates of coactivators and TFs can form at enhancers in cells despite being present at concentrations that are too low for phase separation *in vitro*. We also found that at these low protein concentrations, DNA elements with TF binding sites serve as a scaffold for the phase separated transcriptional condensate. However, at high protein concentrations, the DNA element acts as a nucleation seed only, and is not necessary for condensate stability. This result suggests an explanation for why pathological condensate behavior is often linked to over-expression of coactivators (Zhu et al., 1999).

By considering the competing thermodynamic forces of entropy loss and energy gain that control phase separation, we described how a combination of specific TF-DNA interactions and weak cooperative interactions between IDRs of TFs and coactivators are required for transcriptional condensate formation. These parameters must be above sharply defined thresholds for phase separation to occur. The necessary sharp threshold for TF-DNA interactions results in formation of stable transcriptional condensates at specific genomic loci containing cognate TF binding sites. That there is a threshold affinity between IDRs of the interacting species for condensate formation implies that molecules with IDRs with certain complementary characteristics, such as those contained in transcriptional molecules like TFs and coactivators, will be incorporated in the condensate. Note, however, that IDR-IDR interactions are not of the “lock-key” type, and similar cooperative interactions between a common set of IDRs can mediate transcriptional condensate formation at different genomic loci.

Importantly, we find that DNA motifs with dense clusters of TF binding sites that exceed a sharply defined valency threshold promote transcriptional condensate formation, and the same findings are mirrored for enhancer activity in cells. Bioinformatic analyses reveal that these features also characterize enhancer regions in mammalian genomes, and increases in the recruitment of transcriptional molecules at different loci are correlated in a highly non-linear way with motif density.

Taken together our results suggest the following model for a general cooperative mechanism that contributes to assembling the transcriptional machinery at enhancers, perhaps especially at super-enhancers. Dense clusters of a particular TF’s binding sites, with the valency of binding sites exceeding a sharply defined threshold, drive localized formation of transcriptional condensates at a specific genomic locus. The condensate, which recruits and stabilizes various transcriptional molecules, forms by the universal cooperative mechanism of phase separation. Condensate formation requires weak cooperative interactions between the IDRs of TFs and coactivators (Figure 3). Although different molecular grammars may describe different types of IDR-IDR interactions, these interactions are relatively non-specific, and the same coactivator IDRs can assemble within condensates at different enhancers.

This model is consistent with the observation that clusters of TF binding sites in specific loci in the genome can often correctly predict active enhancers (Berman et al., 2002; Markstein et al., 2002; Rajewsky et al., 2002). But, at the same time, the model suggests that clusters of TF binding sites do not always code for active enhancers (Mansour et al., 2014) because they do not contain a sufficiently high valency of TF binding sites. According to our model, the threshold valency required for condensate formation depends upon the strength of TF-DNA interactions as well as IDR-IDR interactions between transcriptional molecules.

Our model can also describe situations where insertion of a relatively small DNA element that binds to a master TF that regulates cell type specific gene expression programs can stabilize TFs that bind weakly to adjacent binding sites, and recruit the transcriptional machinery in condensates. We carried out simulations with a DNA sequence comprised of two types of binding sites – those that bind strongly to a TF and others that bind weakly. As Figure S6A shows, a transcriptional condensate forms at such a locus beyond a threshold fraction of high-affinity (master) TF binding sites. This is because the cooperative process of condensate formation recruits and stabilizes the transcriptional machinery once the valency of strong TF binding sites exceeds a certain value. This result may explain why a relatively small insertion of a TF binding site into a region that contained an inactive cluster of binding sites for other TFs, resulted in the formation of a super-enhancer in T-ALL cells (Mansour et al., 2014).

While our model explicitly incorporates enhancer DNA, TFs, and coactivators, the underlying mechanistic framework can be extended to understand diverse condensates whose constituents are characterized by a mix of highly specific and weak cooperative interactions. Examples may include condensates in miRNA-mediated silencing (Sheu-Gruttadauria and MacRae, 2018), RNA splicing (Wang et al., 2018), heterochromatin-organization (Larson et al., 2017; Strom et al., 2017), histone locus body assembly (Nizami et al., 2010), lncRNA-mediated paraspeckle formation (Fox et al., 2018; Yamazaki et al., 2018), and in polycomb-mediated transcriptional silencing (Tatavosian et al., 2018).

An implication of our model is that features encoded in DNA can locally condense transcription machinery, without the need for globally tuning protein production or activity. A recent study mapping the phase diagram of engineered proteins in cells also arrives at similar conclusions (Bracha et al., 2018), suggesting a controllable way to spatiotemporally organize condensates. Our results also show that, at TF-DNA affinities close to the condensation threshold, condensates dynamically form and disassemble at DNA elements (Figure S6B,C), which may explain observations of transient assemblies of coactivator and RNA Pol II (Cho et al., 2018), and contribute to observed heterogeneities in gene expression (Thattai and van Oudenaarden, 2001). Given our results, future studies of processes like local RNA synthesis and chromatin modifications, which locally modulate valency and specificity of interacting species, should be fruitful.

## Supporting information

## Acknowledgments

We acknowledge members of Chakraborty and Young labs for helpful discussions. We acknowledge the support of Wendy Salmon and the W.M Keck Microscopy Facility at the Whitehead Institute.

## Funding

This work was supported by grants from the NSF (PHY-1743900) (A.K.C., R.A.Y., and P.A.S), the NIH (GM123511) (R.A.Y.), (P01-CA042063) (P.A.S.), and, in part, by a Koch Institute Support (core) grant (P30-CA14051) from the National Cancer Institute (P.A.S). Additional support was provided by: Damon Runyon Cancer Research Foundation Fellowship (2309-17) (B.R.S.), NSF Graduate Research Fellowship (A.V.Z.), NWO Rubicon Fellowship (J.S.), Swedish Research Council Postdoctoral Fellowship (VR 2017-00372) (A.B.), and an NIH training grant (T32CA009172) (I.A.K.).

## Author Contributions

Conceptualization: K.S., B.R.S., P.A.S., A.K.C., R.A.Y. Methodology: K.S., B.R.S., A.K.C., A.V.Z, J.S., E.L.C., I.A.K., A.B. Software: K.S. Formal Analysis: K.S. Investigation: K.S., B.R.S., A.V.Z, J.S., E.L.C., I.A.K., A.B. Resources: J.S., I.A.K., A.B., N.M.H, P.A.S., A.K.C., R.A.Y. Data Curation: K.S. Writing - Original Draft: K.S., B.R.S., P.A.S., A.K.C., R.A.Y. Writing – Reviewing and Editing: All authors. Visualization: K.S., B.R.S., A.V.Z. Supervision: P.A.S., A.K.C., R.A.Y. Funding Acquisition: P.A.S., A.K.C., R.A.Y.

## Declaration of interests

R.A.Y. is a founder and shareholder of Syros Pharmaceuticals, Camp4 Therapeutics, and Omega Therapeutics. I.A.K. is a consultant to InfiniteMD, Best Doctors, and Foundation Medicine, and is a shareholder of InfiniteMD. All other authors declare no competing interests.

## References

Anderson, J.A., Lorenz, C.D., and Travesset, A. (2008). General purpose molecular dynamics simulations fully implemented on graphics processing units. J. Comput. Phys. 227, 5342–5359.

Banani, S.F., Lee, H.O., Hyman, A.A., and Rosen, M.K. (2017). Biomolecular condensates: organizers of cellular biochemistry. Nat. Rev. Mol. Cell Biol. 18, 285–298.

Berman, B.P., Nibu, Y., Pfeiffer, B.D., Tomancak, P., Celniker, S.E., Levine, M., Rubin, G.M., and Eisen, M.B. (2002). Exploiting transcription factor binding site clustering to identify cis-regulatory modules involved in pattern formation in the Drosophila genome. Proc. Natl. Acad. Sci. U. S. A. 99, 757–762.

Boija, A., Klein, I.A., Sabari, B.R., Dall’Agnese, A., Coffey, E.L., Zamudio, A. V., Li, C.H., Shrinivas, K., Manteiga, J.C., Hannett, N.M., et al. (2018). Transcription Factors Activate Genes through the Phase-Separation Capacity of Their Activation Domains. Cell.

Bracha, D., Walls, M.T., Wei, M.-T., Zhu, L., Kurian, M., Avalos, J.L., Toettcher, J.E., and Brangwynne, C.P. (2018). Mapping Local and Global Liquid Phase Behavior in Living Cells Using Photo-Oligomerizable Seeds. Cell 175, 1467–1480.e13.

Brady, J.P., Farber, P.J., Sekhar, A., Lin, Y.-H., Huang, R., Bah, A., Nott, T.J., Chan, H.S., Baldwin, A.J., Forman-Kay, J.D., et al. (2017). Structural and hydrodynamic properties of an intrinsically disordered region of a germ cell-specific protein on phase separation. Proc. Natl. Acad. Sci. U. S. A. 114, E8194–E8203.

Brangwynne, C.P., Tompa, P., and Pappu, R. V. (2015). Polymer physics of intracellular phase transitions. Nat. Phys. 11, 899–904.

Cho, W.-K., Spille, J.-H., Hecht, M., Lee, C., Li, C., Grube, V., and Cisse, I.I. (2018). Mediator and RNA polymerase II clusters associate in transcription-dependent condensates. Science 361, 412–415.

Chong, S., Dugast-Darzacq, C., Liu, Z., Dong, P., Dailey, G.M., Cattoglio, C., Heckert, A., Banala, S., Lavis, L., Darzacq, X., et al. (2018). Imaging dynamic and selective low-complexity domain interactions that control gene transcription. Science 361.

Fox, A.H., Nakagawa, S., Hirose, T., and Bond, C.S. (2018). Paraspeckles: Where Long Noncoding RNA Meets Phase Separation. Trends Biochem. Sci. 43, 124–135.

Fukaya, T., Lim, B., and Levine, M. (2016). Enhancer Control of Transcriptional Bursting. Cell 166, 358–368.

Hnisz, D., Shrinivas, K., Young, R.A., Chakraborty, A.K., and Sharp, P.A. (2017). A Phase Separation Model for Transcriptional Control. Cell 169, 13–23.

Jolma, A., Yin, Y., Nitta, K.R., Dave, K., Popov, A., Taipale, M., Enge, M., Kivioja, T., Morgunova, E., and Taipale, J. (2015). DNA-dependent formation of transcription factor pairs alters their binding specificity. Nature 527, 384–388.

Jung, C., Bandilla, P., von Reutern, M., Schnepf, M., Rieder, S., Unnerstall, U., and Gaul, U. (2018). True equilibrium measurement of transcription factor-DNA binding affinities using automated polarization microscopy. Nat. Commun. 9, 1605.

Lambert, S.A., Jolma, A., Campitelli, L.F., Das, P.K., Yin, Y., Albu, M., Chen, X., Taipale, J., Hughes, T.R., and Weirauch, M.T. (2018). The Human Transcription Factors. Cell 172, 650–665.

Larson, A.G., Elnatan, D., Keenen, M.M., Trnka, M.J., Johnston, J.B., Burlingame, A.L., Agard, D.A., Redding, S., and Narlikar, G.J. (2017). Liquid droplet formation by HP1α suggests a role for phase separation in heterochromatin. Nature 236–240.

Levo, M., and Segal, E. (2014). In pursuit of design principles of regulatory sequences. Nat. Rev. Genet. 15, 453–468.

Long, H.K., Prescott, S.L., and Wysocka, J. (2016). Ever-Changing Landscapes: Transcriptional Enhancers in Development and Evolution. Cell 167, 1170–1187.

Lu, H., Yu, D., Hansen, A.S., Ganguly, S., Liu, R., Heckert, A., Darzacq, X., and Zhou, Q. (2018). Phase-separation mechanism for C-terminal hyperphosphorylation of RNA polymerase II. Nature 558, 318–323.

Maniatis, T., Falvo, J. V, Kim, T.H., Kim, T.K., Lin, C.H., Parekh, B.S., and Wathelet, M.G. (1998). Structure and function of the interferon-beta enhanceosome. Cold Spring Harb. Symp. Quant. Biol. 63, 609–620.

Markstein, M., Markstein, P., Markstein, V., and Levine, M.S. (2002). Genome-wide analysis of clustered Dorsal binding sites identifies putative target genes in the Drosophila embryo. Proc. Natl. Acad. Sci. 99, 763–768.

Morgunova, E., and Taipale, J. (2017). Structural perspective of cooperative transcription factor binding. Curr. Opin. Struct. Biol. 47, 1–8.

Nizami, Z., Deryusheva, S., and Gall, J.G. (2010). The Cajal Body and Histone Locus Body. Cold Spring Harb. Perspect. Biol. 2, a000653.

Nott, T.J., Petsalaki, E., Farber, P., Jervis, D., Fussner, E., Plochowietz, A., Craggs, T.D., Bazett-Jones, D.P., Pawson, T., Forman-Kay, J.D., et al. (2015). Phase Transition of a Disordered Nuage Protein Generates Environmentally Responsive Membraneless Organelles. Mol. Cell 57, 936–947.

Ptashne, M., and Gann, A. (1997). Transcriptional activation by recruitment. Nature 386, 569–577.

Rajewsky, N., Vergassola, M., Gaul, U., and Siggia, E.D. (2002). Computational detection of genomic cis-regulatory modules applied to body patterning in the early Drosophila embryo. BMC Bioinformatics 3, 30.

Rowley, M.J., Nichols, M.H., Lyu, X., Ando-Kuri, M., Rivera, I.S.M., Hermetz, K., Wang, P., Ruan, Y., and Corces, V.G. (2017). Evolutionarily Conserved Principles Predict 3D Chromatin Organization. Mol. Cell 67, 837–852.e7.

Sabari, B.R., Dall’Agnese, A., Boija, A., Klein, I.A., Coffey, E.L., Shrinivas, K., Abraham, B.J., Hannett, N.M., Zamudio, A. V., Manteiga, J.C., et al. (2018). Coactivator condensation at super-enhancers links phase separation and gene control. Science 361.

Schwarzer, W., Abdennur, N., Goloborodko, A., Pekowska, A., Fudenberg, G., Loe-Mie, Y., Fonseca, N.A., Huber, W., Haering, C., Mirny, L., et al. (2017). Two independent modes of chromatin organization revealed by cohesin removal. Nature 551, 51.

Sheu-Gruttadauria, J., and MacRae, I.J. (2018). Phase Transitions in the Assembly and Function of Human miRISC. Cell 173, 946–957.e16.

Shin, Y., and Brangwynne, C.P. (2017). Liquid phase condensation in cell physiology and disease. Science 357.

Shlyueva, D., Stampfel, G., and Stark, A. (2014). Transcriptional enhancers: from properties to genome-wide predictions. Nat. Rev. Genet. 15, 272–286.

Slattery, M., Zhou, T., Yang, L., Dantas Machado, A.C., Gordân, R., and Rohs, R. (2014). Absence of a simple code: how transcription factors read the genome. Trends Biochem. Sci. 39, 381–399.

Spitz, F., and Furlong, E.E.M. (2012). Transcription factors: from enhancer binding to developmental control. Nat. Rev. Genet. 13, 613–626.

Strom, A.R., Emelyanov, A. V., Mir, M., Fyodorov, D. V., Darzacq, X., and Karpen, G.H. (2017). Phase separation drives heterochromatin domain formation. Nature 547, 241–245.

Tatavosian, R., Kent, S., Brown, K., Yao, T., Duc, H.N., Huynh, T.N., Zhen, C.Y., Ma, B., Wang, H., and Ren, X. (2018). Nuclear condensates of the Polycomb protein chromobox 2 (CBX2) assemble through phase separation. J. Biol. Chem. jbc.RA118.006620.

Thattai, M., and van Oudenaarden, A. (2001). Intrinsic noise in gene regulatory networks. Proc. Natl. Acad. Sci. 98, 8614–8619.

Tsai, A., Muthusamy, A.K., Alves, M.R., Lavis, L.D., Singer, R.H., Stern, D.L., and Crocker, J. (2017). Nuclear microenvironments modulate transcription from low-affinity enhancers. Elife 6.

Wang, A., Conicella, A.E., Schmidt, H.B., Martin, E.W., Rhoads, S.N., Reeb, A.N., Nourse, A., Ramirez Montero, D., Ryan, V.H., Rohatgi, R., et al. (2018). A single N-terminal phosphomimic disrupts TDP-43 polymerization, phase separation, and RNA splicing. EMBO J. 37, e97452.

Wasserman, W.W., and Sandelin, A. (2004). Applied bioinformatics for the identification of regulatory elements. Nat. Rev. Genet. 5, 276–287.

Wei, M.-T., Elbaum-Garfinkle, S., Holehouse, A.S., Chen, C.C.-H., Feric, M., Arnold, C.B., Priestley, R.D., Pappu, R. V., Brangwynne, C.P., Chih, C., et al. (2017). Phase behaviour of disordered proteins underlying low density and high permeability of liquid organelles. Nat. Chem. 9, 1118–1125.

Weirauch, M.T., Yang, A., Albu, M., Cote, A.G., Montenegro-Montero, A., Drewe, P., Najafabadi, H.S., Lambert, S.A., Mann, I., Cook, K., et al. (2014). Determination and inference of eukaryotic transcription factor sequence specificity. Cell 158, 1431–1443.

Whyte, W.A., Orlando, D.A., Hnisz, D., Abraham, B.J., Lin, C.Y., Kagey, M.H., Rahl, P.B., Lee, T.I., and Young, R.A. (2013). Master Transcription Factors and Mediator Establish Super-Enhancers at Key Cell Identity Genes. Cell 153, 307–319.

Wunderlich, Z., and Mirny, L.A. (2009). Different gene regulation strategies revealed by analysis of binding motifs. Trends Genet. 25, 434–440.

Yamazaki, T., Souquere, S., Chujo, T., Kobelke, S., Chong, Y.S., Fox, A.H., Bond, C.S., Nakagawa, S., Pierron, G., and Hirose, T. (2018). Functional Domains of NEAT1 Architectural lncRNA Induce Paraspeckle Assembly through Phase Separation. Mol. Cell 70, 1038–1053.e7.

Zhu, Y., Qi, C., Jain, S., Le Beau, M.M., Espinosa, R., Atkins, G.B., Lazar, M.A., Yeldandi, A. V, Rao, M.S., and Reddy, J.K. (1999). Amplification and overexpression of peroxisome proliferator-activated receptor binding protein (PBP/PPARBP) gene in breast cancer. Proc. Natl. Acad. Sci. U. S. A. 96, 10848–10853.

