## Supplementary material for "Enhancer features that drive formation of transcriptional condensates"

[10]Co-first authors

**This PDF file includes:**

Methods
Supplementary References
Figs. S1-S6
Tables S1-S3
Caption for Movies S1-S2

#### METHODS

##### Cells

V6.5 murine embryonic stem cells were a gift from R. Jaenisch of the Whitehead Institute. V6.5 are male cells derived from a C57BL/6(F) x 129/sv(M) cross.

##### Cell culture conditions

V6.5 murine embryonic stem cells were grown in 2i + LIF conditions on 0.2% gelatinized (Sigma, G1890) tissue culture plates. 2i + LIF media contains the following: 967.5 mL DMEM/F12 (GIBCO 11320), 5 mL N2 supplement (GIBCO 17502048), 10 mL B27 supplement (GIBCO 17504044), 0.5mML-glutaminae (GIBCO 25030), 0.5X non-essential amino acids (GIBCO 11140), 100 U/mL Penicillin-Streptomycin (GIBCO 15140), 0.1 mM b-mercaptoethanol (Sigma), 1 uM PD0325901 (Stemgent 04- 0006), 3 uM CHIR99021 (Stemgent 04-0004), and 1000 U/mL recombinant LIF (ESGRO ESG1107). Cells were negative for mycoplasma.

**Developing coarse-grained simulations of DNA, TFs, and coactivators**

We set up a coarse-grained molecular-dynamics simulation to model 3 different components – TFs, DNA, and coactivators, employing the HOOMD simulation framework (Anderson et al., 2008; Glaser et al., 2015). Briefly, the DNA chain was modeled as beads on a string, with two types of monomers. "Active" DNA units were modeled by tessellating a sphere ($diameter = 1/3\ unit$), using the rigid-body feature(Nguyen et al., 2011), to form a roughly cubical monomer of unit side length (Fig 1A). Binding patches were modeled as rigid particles along the cubic face centers, with as many patches added as number of binding sites per monomer. Tessellation of active DNA monomers enabled 1:1 binding interactions, facilitated by excluded volume interactions from other tessellated spheres. "Inactive" DNA monomers were modeled as spherical monomers of unit diameter without any binding patches. TFs and coactivators were modeled employing two different methods – explicit-IDR (Fig 1A) and implicit-IDR models (Fig S3A). In the explicit-IDR framework, TFs and coactivators were designed in a modular fashion. The "structured" domain was modeled as a spherical monomer of diameter $d = 0.75$ units.s. IDRs were constructed by tethering a polymeric tail to the spherical domain, with TFs having shorter chains (4 monomers of $d = 1/3\ unit$) than coactivators (9 monomers of $d = 1/3\ unit$), to mimic the differential size of disordered regions. In the implicit-IDR model, TFs and coactivators were modeled as spherical monomers of unit diameter. All monomers had the same density. In both methods, DNA binding patches on proteins were modeled as rigid particles buried in the "structured" domains.

Non-bonding interactions between any two particles (including binding patches) were modeled using a truncated, shifted, and size-normalized LJ potential (U) with hard-core repulsion (particles don't overlap), derived in the following form:

$$U_{ij}(\vec{r}) = \begin{pmatrix} P_{ij}(r) - P_{ij}(r^*) & r \leq r^* \\ 0 & r > r^* \end{pmatrix}$$

$$P_{ij}(r) = 4\epsilon_{ij} \times ((^{\sigma}/_{r})^{12} - (^{\sigma}/_{r})^{6})$$

$$\sigma = 0.5 \times \left(d_i + d_j\right), r^* = 2.5 \times \sigma$$

Bonding interactions between neighboring monomers on a chain were modeled using a harmonic potential with hard-cores, with a spring constant $k = 1e4$. All energy units are scaled to kT units, with kT=1.

The strength of various interactions was set based on the rationale stated in main text. Typical TF-DNA binding affinities are strong and in the range of nanomolar (Jung et al., 2018) disassociation constants i.e. $K_D \approx 10^{-9}M$. The Gibbs free enthalpy change of binding can be approximately calculated as $\Delta G \approx -kTln(K_D) \approx 20kT$. Thus, specific monovalent DNA interactions were set to high affinities - for e.g. $\epsilon_{DNA-TF} = 20kT$ in fig 1B,2A, $\epsilon_{DNA-TF} = 16\ kT$ in fig S3A-B. IDR interactions were much weaker and individual interactions are often of the order of thermal fluctuations (Brady et al., 2017; Nott et al., 2015; Wei et al., 2017) i.e. order kT. Thus, we set $\epsilon_{IDR} \sim kT$ between monomers on the IDR chain. For the implicit-IDR model reported in Fig S3, multivalent interactions were approximated by a weak LJ potential between particles, for e.g., $\epsilon_{TF-coA} = 1.5kT, \epsilon_{coA-coA} = 1.5kT, \epsilon_{TF-TF} = 1.0\ kT$ . The key qualitative results i.e multivalent DNA acts as scaffold for phase separation at low protein concentrations and seed at higher protein levels, has been reproduced for different choices of interaction parameters guided by the rationale above.

Particles are randomly initialized in the periodic simulation box, and randomly re-seeded for each replicate trajectory, with the Langevin thermostat. Friction coefficients were $\gamma = 1$ for proteins and $\gamma = 100$ for DNA, to mimic chromosomal motion damping. Initial velocities were drawn from the Boltzmann distribution. First, simulations were run with small time steps ($dt = 5 \times 10^{-6}$) to prevent randomly generated "high-energy" configurations from blowing up and to relax the system to the thermostat temperature. These "warm-up" period ($t \sim 0.1\ units$) is much smaller than the time to reach steady-state $t_{ss} \sim (1000\ units)$, so these warm-up data points are not used in any analysis. All simulations are run with a single DNA chain.

Explicit-IDR simulations are run for at least $25e6$ steps to accurately recapitulate dynamics and reach steady-state, whilst implicit-IDR simulations are run for $5e6$ steps. The slowing down of explicit-IDR simulations (due to slower explicit-IDR dynamics), combined with additional pair-wise interaction computations (explicit pairwise calls for all monomers, which are an order of magnitude more particles for explicit-IDR simulations, scale as $\sim N^2$ for N monomers), cause computation times for single trajectories to be ~50-100 times longer than the implicit-IDR version. Trajectory states were logged in the highly compressed, binarized GSD format every $50000$ steps, while observables were logged every 20000 steps.

To probe the role of DNA in our simulations, after steady-state is reached, interactions with the DNA binding sites are switched off. Interactions are switched off by replacing all binding patches with "ghost" patches, with no energetic benefits. Simulations are typically continued for the same amount of steps before disrupting TF-DNA interactions to accurately sample steady-state. A brief overview of key parameters used in main/supplementary figures is found below in Table S1. The MD code for running analysis will be made freely available upon publication.

**Analysis of simulation data**

Broadly, analyses of simulation data were split into on-the-fly calculations employing the Freud package (https://freud.readthedocs.io/en/stable/installation.html), as well as post-simulation calculations that leverage a combination of various libraries which interface with python – including *numpy*, *scipy*, *freud*, *matplotlib*, and *fresnel*. On-the-fly calculations include:

1. In-built functions for logging potential energy, kinetic energy, and temperature.

2. Number of monomers in largest cluster and radius of largest cluster: A call-back routine was implemented that used Freud to estimate the size of the largest connected cluster with $r = 1.4d_{max}$ ($d_{max}$ diameter of largest monomer) to identify largest cluster. This largest cluster size is relatively insensitive to studied choices of parameter $r = 1.25, 1.35, 1.45\ d_{max}$. Every reported plot with scaled size at steady state, which is the number of molecules in the largest cluster divided by number of binding sites (Fig 1B), reports the mean in the dark line, and one standard deviation in the shaded background.

For post-simulation calculations, data was read from GSD formats using the gsd module. Explicit-IDR simulation trajectory data was parsed to convert from number of molecules to number of chains, while following the other steps as mentioned above.

The entropy was calculated in Fig 2 and Fig S3 by identifying the number of molecules in the largest cluster (in the case of the explicit-IDR simulations, each polymer was counted as one molecule), and adding a value of $kTln(\frac{4/3\pi R_g^3}{V_{free}})$ for each molecule in the condensed phase. $V_{free}$ was computed as the total volume minus the excluded volume occupied by all molecules. For the fluctuation analysis in Fig S6B, the variance in largest cluster size of individual stochastic trajectories was computed and averaged at steady state. This value was normalized by the scaled cluster size, to compute the scaling of fluctuations beyond the usual $\sqrt{N}$ finite-size effects.

**Hi-C analysis of simulation data**

For Hi-C maps represented in Fig 5B, the following analysis protocol was employed. After individual trajectories reached steady-state, the position of each DNA monomer along the chain was logged at every time step. Monomers closer than ($r = 3.0$ units) a distance at a time $t$ are "cross-linked" i.e. they count as an interacting pair. The qualitative interaction maps reported in Fig 5B are robust to other tested values of crosslinking radius in the regime of $2.5 < r < 4$ units. The pairwise contact frequency matrix is then constructed by averaging over interactions over a time window at steady state per trajectory, as well as averaging over 10 replicate trajectories per simulation condition. The contact matrix is visualized using the *seaborn* and *matplotlib* packages in python3.

**Visualization of simulation data:**

All simulation data-sets were analyzed in python, with the aid of *matplotlib*, to generate publication-ready figures. Simulation movies were generated by stitching together down-sampled frames (once every 100000 steps) of individual stochastic trajectories, using Fresnel to render scenes with the same color palette used in Fig 1A, and PIL to store image arrays as gifs. After storing the gifs, these files were converted to .mp4 movies externally and subs are added at the frame at which TF-DNA interactions are turned off.

**Quantitative immunoblot**

Determination of number of MED1 molecules per cell and concentration by linear regression analysis. Quantitative Western Blotting was carried out as described in (Lin et al., 2012). Cell number was determined using a Countless II FL Automated Cell Counter (Thermo Fisher Scientific). Cells were lysed with Cell Lytic M (Sigma) with protease inhibitors at various concentrations and denatured in DTT and XT Sample Buffer (Biorad) at 90°C for 5 minutes. Purified recombinant MED1-IDR was used as a standard and loaded in the amounts depicted in the figure in the same gel as the cell lysates. Lysates and standards were run on a 3%–8% Tris-acetate gel at 80 V for ~2 hrs, followed by 120 V until dye front reached the end of the gel. Protein was then wet transferred to a 0.45 μm PVDF membrane (Millipore, IPVH00010) in ice-cold transfer buffer (25 mM Tris, 192 mM glycine, 10% methanol) at 300 mA for 2 hours at 4°C. After transfer the membrane was blocked with 5% non-fat milk in TBS for 1 hour at room temperature, shaking. Membrane was then incubated with 1:1,000 anti-MED1 (Assay Biotech B0556) diluted in 5% non-fat milk in TBST and incubated overnight at 4°C, with shaking. The membrane was then washed three times with TBST for 5 minutes at room temperature shaking for each wash. Membrane was incubated with 1:10,000 secondary antibody conjugated to HRP for 1 hr at RT and washed three times in TBST for 5 minutes. Membranes were developed with ECL substrate (Thermo Scientific, 34080) and imaged using a CCD camera or exposed using film. Band intensities were determined using ImageJ. Number of molecules per cell was determined by linear regression analysis through the origin using Prism 7. The concentration of MED1 was calculated using nuclear volumes obtained by analysis of Hoechst (Life Technologies)-stained mouse embryonic stem cells in ImageJ and assuming all MED1 molecules reside in the nucleus.

**Protein purification**

Proteins were purified as in (Boija et al., 2018; Sabari et al., 2018). cDNA encoding the genes of interest or their IDRs were cloned into a modified version of a T7 pET expression vector. The base vector was engineered to include a 5’ 6xHIS followed by either mEGFP or mCherry and a 14 amino acid linker sequence “GAPGSAGSAAGGSG.” NEBuilder® HiFi DNA Assembly Master Mix (NEB E2621S) was used to insert these sequences (generated by PCR) in-frame with the linker amino acids. Vectors expressing mEGFP or mCherry alone contain the linker sequence followed by a STOP codon. Mutant sequences were synthesized as gBlocks (IDT) and inserted into the same base vector as described above. All expression constructs were sequenced to ensure sequence identity. For protein expression plasmids were transformed into LOBSTR cells (gift of Chessman Lab) and grown as follows. A fresh bacterial colony was inoculated into LB media containing kanamycin and chloramphenicol and grown overnight at 37°C. Cells containing the MED1-IDR constructs were diluted 1:30 in 500ml room temperature LB with freshly added kanamycin and chloramphenicol and grown 1.5 hours at 16°C. IPTG was added to 1mM and growth continued for 18 hours. Cells were collected and stored frozen at -80°C. Cells containing all other constructs were treated in a similar manner except they were grown for 5 hours at 37°C after IPTG induction. 500ml cell pellets were resuspended in 15ml of Buffer A

(50mM Tris pH7.5, 500 mM NaCl) containing 10mM imidazole and cOmplete protease inhibitors, sonicated, lysates cleared by centrifugation at 12,000g for 30 minutes at 4°C, added to 1ml of pre-equilibrated Ni-NTA agarose, and rotated at 4°C for 1.5 hours. The slurry was poured into a column, washed with 15 volumes of Buffer A containing 10mM imidazole and protein was eluted 2 X with Buffer A containing 50mM imidazole, 2 X with Buffer A containing 100mM imidazole, and 3 X with Buffer A containing 250mM imidazole.

**Production of fluorescent DNA**

Gene fragments were synthesized by either GeneWiz or IDT and cloned into a pUC19 vector using HiFi Assembly (NEB) so that the sequence was immediately flanked by M13(-21) and M13 reverse primer sequences. 5'-fluorescently labeled (Cy5) M13(-21) (/5Cy5/ TGTAAAACGACGGCCAGT) and M13 reverse (/5Cy5/ CAGGAAACAGCTATGAC) primers (IDT) were used to PCR amplify the synthetic DNA sequence, yielding a fluorescently labeled PCR product. Fluorescent PCR products were gel-purified (Qiagen) and eluted products were further purified using NEB Monarch PCR purification to remove any residual contaminants. The octamer motif sequence "ATTTGCAT" from the immunoglobulin kappa promoter was used as the TF binding site. All PCR products used are 377 bp. The sequences of PCR products are provided in Table S2.

**In vitro droplet assay**

Recombinant GFP or mCherry fusion proteins were concentrated and desalted to an appropriate protein concentration and 125mM NaCl using Amicon Ultra centrifugal filters (30K MWCO, Millipore) in Buffer D(125) (50mM Tris-HCl pH 7.5, 10% glycerol, 1mM DTT). Fluorescent PCR products were concentration normalized in Buffer D(0) (50mM Tris-HCl pH 7.5, 10% glycerol, 1mM DTT). For all droplet assays, DNA was included at 50nM, mEGFP-OCT4 or mEGFP as 1250nM, and mCherry-MED1-IDR at the indicated concentration. Recombinant proteins and DNA were mixed with 10% PEG-8000 as a crowding agent. The final buffer conditions were 50mM Tris-HCl pH 7.5, 100mM NaCl, 10% glycerol, 1mM DTT. The solution was immediately loaded onto a homemade chamber comprising a glass slide with a coverslip attached by two parallel strips of double-sided tape. Slides were then imaged with an Andor confocal microscope with a 150x objective. Unless indicated, images presented are of droplets settled on the glass coverslip. For DNase I experiment, MED1-IDR droplets were allowed to form at indicated concentration in the presence of OCT4 (1250nM) and DNA (50nM) for 5 minutes. To experimental sample, DNase I (Promega RQ1 M6101) was added with manufacturer provided reaction buffer. To control samples, manufacturer provided reaction buffer was added. Both were incubated at 30° C for 10 minutes and subsequently imaged as described above.

**Image analysis for reconstructing experimental phase curves**

A custom analysis pipeline was developed in MATLAB$^{TM}$, building on code described in (Boija et al., 2018). Briefly, droplets were identified by employing a two-step thresholding procedure. First, the image was segmented in the MED1-IDR channel with an intensity threshold ($I_{pixel} > \mu^* + 3\sigma$, where $\mu^*$ is the most probable intensity, representative of background, and $\sigma$ is the width of the distribution) to identify bright pixels. Subsequently, the identified bright pixels were labeled as "condensed" droplet phase after enforcing a minimum droplet size of 9 pixels i.e. atleast 9 clustered pixels had to simultaneously pass the intensity threshold to belong to the condensed phase. In the absence of phase separation, no pixels are identified as belonging to the condensed phase.

For each image, the total intensity in the condensed droplet phase was summed in each channel ($I_{channel,droplet}$), as well as the total background intensity outside droplets ($I_{channel,bulk}$). The condensed fraction in each channel was defined as :

$$c.f._{channel} = \frac{I_{channel,droplet}}{I_{channel,droplet} + I_{channel,bulk}}.$$

The condensed fraction was averaged over replicate images (~ 10 per condition). At very low concentrations or in the absence of observable phase separation, c.f. is close to 0. We repeated the c.f. analysis with different intensity thresholds ($I > \mu^* + 2.5\sigma, I > \mu^* + 3.5\sigma, I > \mu^* + 4\sigma$) and size thresholds ($9,16,25\ pixels$). The qualitative results reported in Fig 1,3,4 and supplementary Fig 4 did not change under these tested conditions.

In all plots of the c.f., solid lines represent the mean condensed fraction and shaded background refers to values one standard deviation above and below the mean. Plots of the condensed fraction were generated by using the *matplotlib* library in python3. In all plots in the main figures (Figs 1F, 3D, 3H, 4D), the condensed fraction in the MED1-IDR channel is reported.

**DNase I image analysis**

Building on the above-described analysis, for each condition, the partition ratio for each replicate image was calculated as $p_{channel} = \frac{<Intensity>_{droplet}}{<Intensity>_{bulk}}$ in various channels. The partition ratio is a proxy for the relative enrichment of molecules in the condensed phase over the bulk phase. Subsequently, the partition ratios for control (without DNAseI) and DNAseI experiments were normalized to the mean partition ratio for the control at same concentration of MED1-IDR. Box-plots were generated using the distribution of normalized partition ratios in the 561(MED1-IDR) channel for Fig. 2B, and in the 640 channel (DNA) for Fig S1B, using PRISM.

**Luciferase reporter assays**

For enhancer activity reporter assays, synthetic enhancer DNA sequences with varying valencies or densities of OCT4 binding sites (see Table S3) were cloned into a previously characterized pGL3-basic construct containing a minimal OCT4 promoter (pGL3-pOCT4)(Whyte et al., 2013). The synthetic enhancer sequences were cloned into the SalI site of the pGL3-pOct4 vector by HiFi DNA Assembly (NEB E2621) with a SalI digested vector and PCR-amplified insert. All cloned constructs were sequenced to ensure sequence identity. 0.4μg of the pGL3-based enhancer plasmids were used to transfect $1x10^5$ murine ESCs in 24-well plates using Lipofectamine 3000 (Thermo Fisher L3000015) according to the manufacturer's instructions. 0.1μg of the pRL-SV40 plasmid was co-transfected in each condition as a luminescence control. Transfected cells were harvested after 24 hours, and luciferase activity was measured using the Dual-Glo Luciferase Assay System (Promega E2920). Luciferase signal was normalized to the signal measured in cells transfected with a construct containing zero OCT4 motifs. Experiments were performed in triplicates.

**Bioinformatic analysis**

Position-weight matrices (PWMS) for *Mus musculus* stem cell master TFs –SOX2 (MA0143), OCT4+SOX2 (MA0142), KLF4 (MA0039), and ESRRB (MA0141), were obtained from the JASPAR database (Khan et al., 2018). 100kb DNA sequences centered on super-enhancers (SEs, N=231), as annotated in (Whyte et al., 2013) were gathered. The same number and length of sequences were randomly subsampled from enhancers (typical enhancers, TEs) annotated in (Whyte et al., 2013), as well as from random genetic loci (Random) on the *mm9* reference genome. FIMO was used to predict individual motif instances in all sequences, against a background uniform random distribution, at a *p-value* threshold of 1e-4.

For the boxplots in Fig 6A, the average motif density is calculated as total number of motifs divided by length of sequence over a 20kb sequence region centered on SEs, TEs, and random loci, normalized in units of motifs/kb. For the line plots in Fig 6B, the whole distribution of motif density is represented along the 100 KB sequence, in bins of 2kb with similar units.

Published ChIP-Seq data-sets are gathered from (Sabari et al., 2018) for MED1, BRD4, RNA Pol II, and input control from GEO: GSE112808. Reads-per-million (rpm) are summed in previously defined regions for SEs, TES, and random using BedTools. For Fig 6C, and supplementary figure S5, the summed rpm values are plotted on a log scale. On the x-axis, the total number of motifs calculated in a 20kb window centered on SEs, TEs, and random loci is plotted. Finally, a linear model is inferred between $\log(ChIP)$ signal and motif values using ordinary least squares regression. The inferred line is plotted in black and 95% confidence intervals are plotted as a shaded gray background. The data is visualized using the *matplotlib* library in python3.

**Statistical analysis for simulation data:**
Steady-state analysis of simulation data-sets in Fig 3 & 4 are reported with solid lines represented by the mean ($\mu$) and fluctuations of individual trajectories in the shaded background, whose boundaries are characterized by one standard deviation away from the mean on either side ($\mu \pm \sigma$). In all figures, the mean represents an average over 10 trajectories.

**Statistical analysis for bioinformatics:**
The inferred linear line in Fig 6C and S6 are generated between the logarithm of the ChIP signal and the motif density, and the $R^2$ reported in the respective captions.

**Statistical analysis for computational image analysis:**
Condensed fraction reported at any given concentration in all figures are averaged over > 10 image-sets, with shaded background representing one standard deviation from the mean condensed fraction.

#### DATA AND SOFTWARE AVAILABILITY

All software and code generated in this project will be available freely upon publication.

**References:**

Anderson, J.A., Lorenz, C.D., and Travesset, A. (2008). General purpose molecular dynamics simulations fully implemented on graphics processing units. J. Comput. Phys. *227*, 5342–5359.

Boija, A., Klein, I.A., Sabari, B.R., Dall'Agnese, A., Coffey, E.L., Zamudio, A. V., Li, C.H., Shrinivas, K., Manteiga, J.C., Hannett, N.M., et al. (2018). Transcription factors activate genes through the phase separation capacity of their activation domains. Cell *in press*.

Brady, J.P., Farber, P.J., Sekhar, A., Lin, Y.-H., Huang, R., Bah, A., Nott, T.J., Chan, H.S., Baldwin, A.J., Forman-Kay, J.D., et al. (2017). Structural and hydrodynamic properties of an intrinsically disordered region of a germ cell-specific protein on phase separation. Proc. Natl. Acad. Sci. U. S. A. *114*, E8194–E8203.

Glaser, J., Nguyen, T.D., Anderson, J.A., Lui, P., Spiga, F., Millan, J.A., Morse, D.C., and Glotzer, S.C. (2015). Strong scaling of general-purpose molecular dynamics simulations on GPUs. Comput. Phys. Commun. *192*, 97–107.

Jung, C., Bandilla, P., von Reutern, M., Schnepf, M., Rieder, S., Unnerstall, U., and Gaul, U. (2018). True equilibrium measurement of transcription factor-DNA binding affinities using automated polarization microscopy. Nat. Commun. *9*, 1605.

Khan, A., Fornes, O., Stigliani, A., Gheorghe, M., Castro-Mondragon, J.A., van der Lee, R., Bessy, A., Chèneby, J., Kulkarni, S.R., Tan, G., et al. (2018). JASPAR 2018: update of the open-access database of transcription factor binding profiles and its web framework. Nucleic Acids

Res. *46*, D260–D266.

Lin, C.Y., Lovén, J., Rahl, P.B., Paranal, R.M., Burge, C.B., Bradner, J.E., Lee, T.I., and Young, R.A. (2012). Transcriptional amplification in tumor cells with elevated c-Myc. Cell *151*, 56–67.

Nguyen, T.D., Phillips, C.L., Anderson, J.A., and Glotzer, S.C. (2011). Rigid body constraints realized in massively-parallel molecular dynamics on graphics processing units. Comput. Phys. Commun. *182*, 2307–2313.

Nott, T.J., Petsalaki, E., Farber, P., Jervis, D., Fussner, E., Plochowietz, A., Craggs, T.D., Bazett-Jones, D.P., Pawson, T., Forman-Kay, J.D., et al. (2015). Phase Transition of a Disordered Nuage Protein Generates Environmentally Responsive Membraneless Organelles. Mol. Cell *57*, 936–947.

Sabari, B.R., Dall'Agnese, A., Boija, A., Klein, I.A., Coffey, E.L., Shrinivas, K., Abraham, B.J., Hannett, N.M., Zamudio, A. V., Manteiga, J.C., et al. (2018). Coactivator condensation at super-enhancers links phase separation and gene control. Science *361*.

Wei, M.-T., Elbaum-Garfinkle, S., Holehouse, A.S., Chih, C., Chen, -Hsiung, Feric, M., Arnold, C.B., Priestley, R.D., Pappu, R. V, and Brangwynne, C.P. (2017). Phase behaviour of disordered proteins underlying low density and high permeability of liquid organelles.

Whyte, W.A., Orlando, D.A., Hnisz, D., Abraham, B.J., Lin, C.Y., Kagey, M.H., Rahl, P.B., Lee, T.I., and Young, R.A. (2013). Master Transcription Factors and Mediator Establish Super-Enhancers at Key Cell Identity Genes.

#### Figure S1

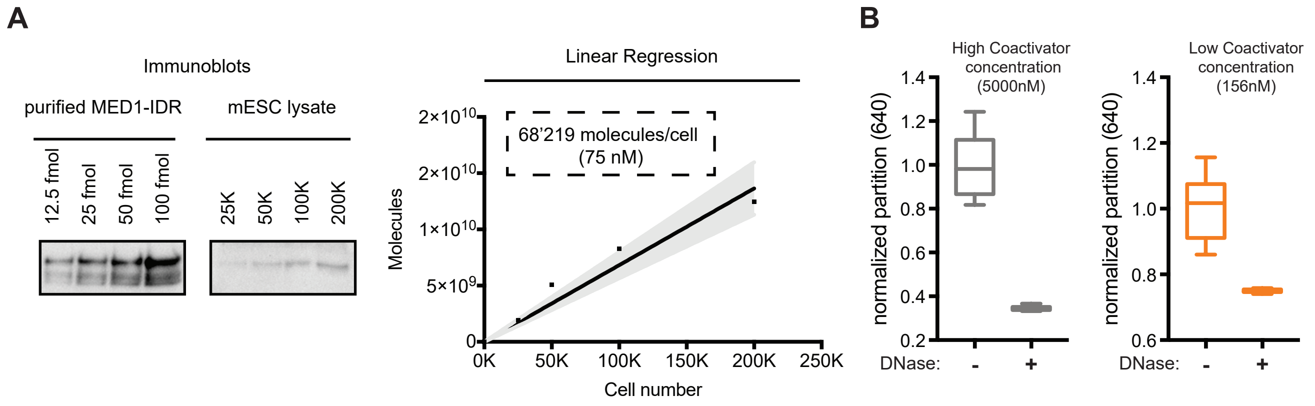


**Figure S1: OCT4-DNA interactions promote phase separation of MED1-IDR at low concentrations, Related to Figures 1,2**

**A.** Immunoblot of recombinant MED1-IDR at indicated concentrations or lysates from the indicated number of cells is shown on the top panel. Linear regression (bottom panel) is carried out to estimate number and concentration of MED1-IDR per cell (dashed box, bottom panel) (see methods for details).

**B.** Box-plot depiction of ODNA_20 partition ratio between condensate and background, at high (gray) and low MED1-IDR concentrations (orange) in conditions without DNase I addition (-) or with DNase I addition (+). The partition ratio is normalized to the (-) condition, showing that addition of DNase I degrades DNA.

#### Figure S2

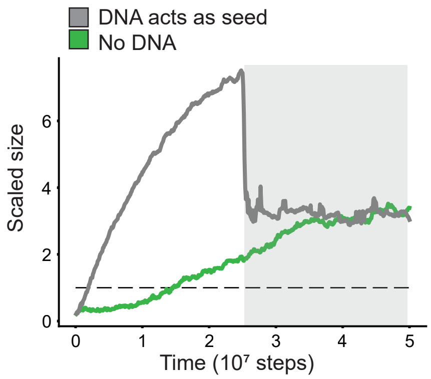


**Figure S2: DNA promotes rate of condensate assembly, but not stability, at high protein concentrations, Related to Figure 2**

Dynamics of condensate assembly at conditions with (grey) and without DNA (green line) is represented by average scaled size on y-axis, and time (in simulation steps after initialization) on the x-axis. DNA promotes rate of assembly at high concentrations. However, DNA is not required for condensate stability, as evidenced by high values of scaled size after disruption of TF-DNA interactions (shown by a dark grey background).

### Figure S3

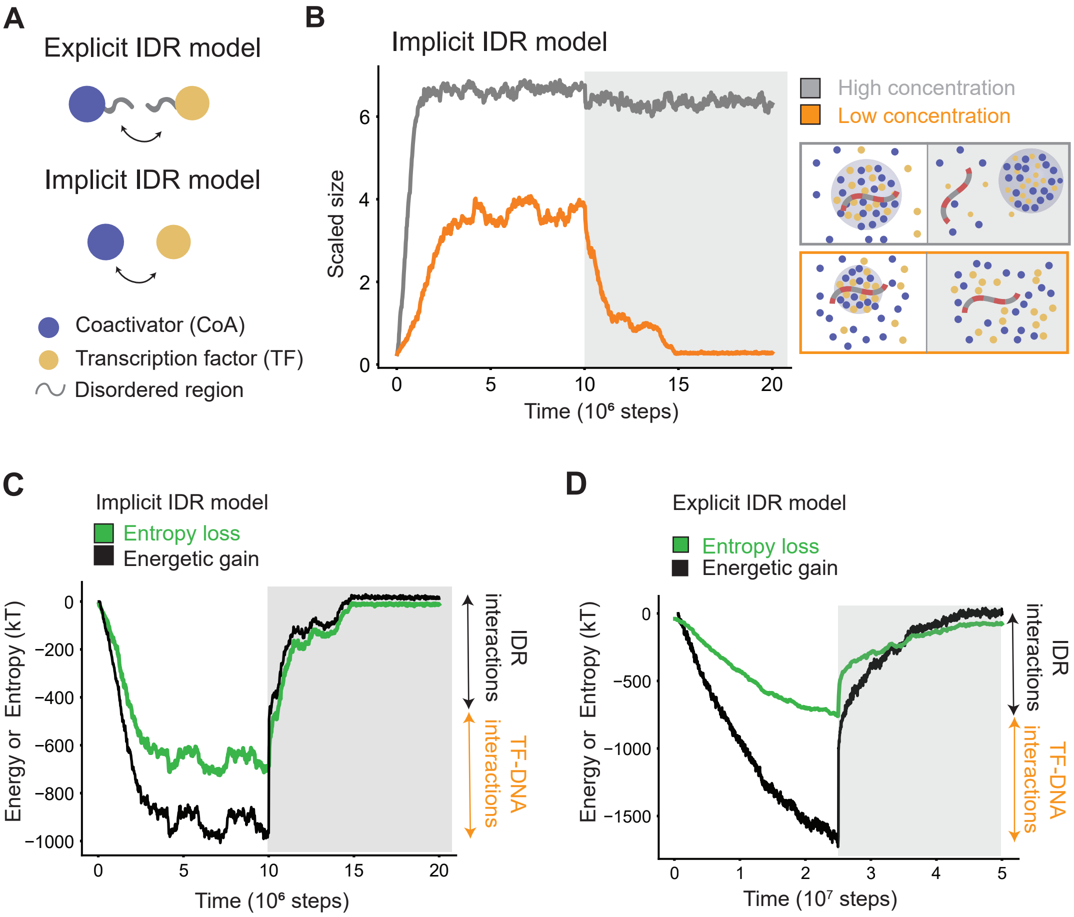


**Figure S3: Simplified computational model recapitulates all features of explicit-IDR model, Related to Figure 2**

**A.** Schematic cartoon of difference between explicit IDR model and implicit IDR model.

**B**. Dynamics of condensate assembly/disassembly at three different protein concentrations (gray = high concentration, orange = low concentration, black = lower concentration) is represented by average scaled size on the y-axis, and time (in simulation steps after initialization) on the x-axis. TF-DNA interactions are disrupted after steady state is reached (shown by a dark gray background). Schematic of phase behavior is presented next to simulation data, enclosed in boxes whose colors match the respective lines.

**C.** Energetic attractions (black line) compensate entropic loss (green line) during condensate assembly, but disruption of TF-DNA interactions (magnitude =orange double arrow) causes dissolution at low concentrations.

**D.** Explicit-IDR simulations show a compensation of entropic loss (green line) by energetic attractions (black line) during condensate assembly, and disruption of TF-DNA interactions causes dissolution. However, the estimate of entropy loss from simulations is an under-count to the total loss of entropy, missing effects of configurational entropy.

#### Figure S4

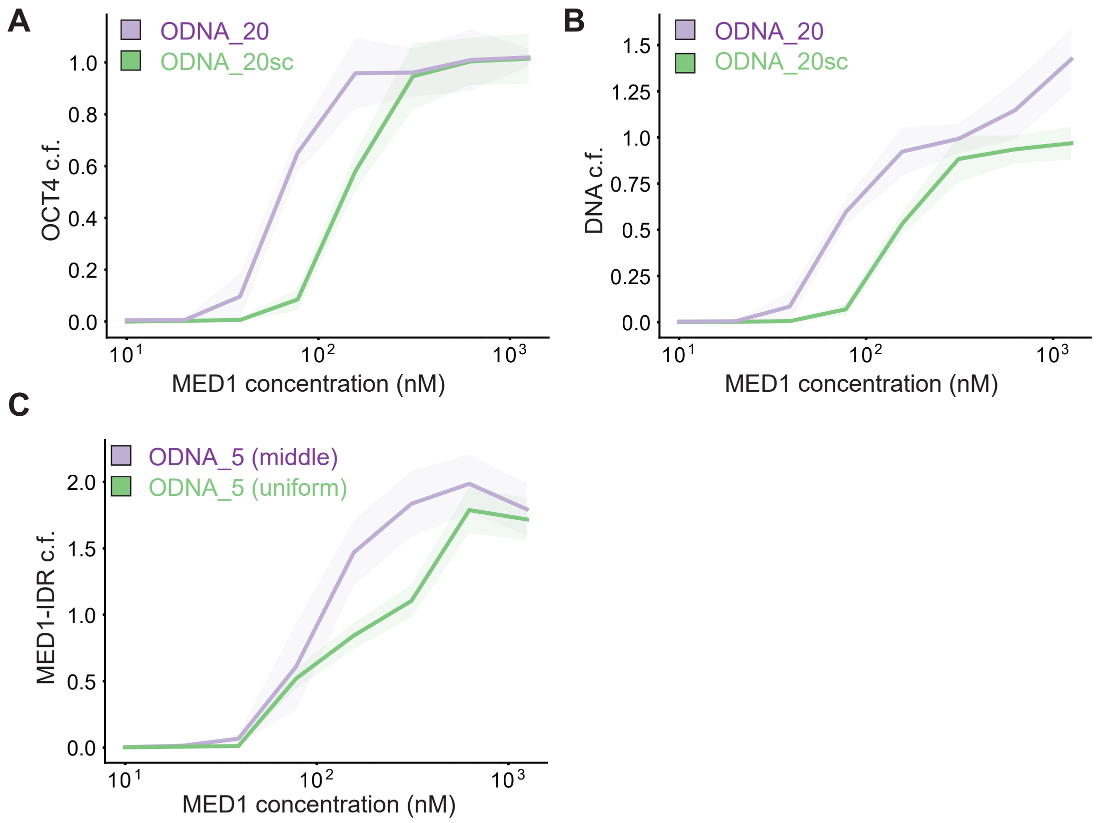


**Figure S4: Phase separation of all components is promoted at lower coactivator concentrations by multivalent DNA with high density of TF binding sites, Related to Figures 3,4**

**A.** Condensed fraction of OCT4 (in units of percentage) for ODNA_20 (purple) and ODNA_20sc (green) across a range of MED1-IDR concentrations.

**B.** Condensed fraction (in units of percentage) of ODNA_20 (purple) and ODNA_20sc (green) across a range of MED1-IDR concentrations.

**C.** Condensed fraction of MED1-IDR (in units of percentage) for ODNA_5_uniform (green) and ODNA_5_middle (purple) across a range of MED1-IDR concentrations. Both DNA sequences have 5 OCT4 binding sites, but ODNA_5_uniform has them spread uniformly, whilst ODNA_5_middle has the binding sites clustered (see Table S2).

Solid lines represent mean and shaded background represents boundaries of mean±std.

#### Figure S5

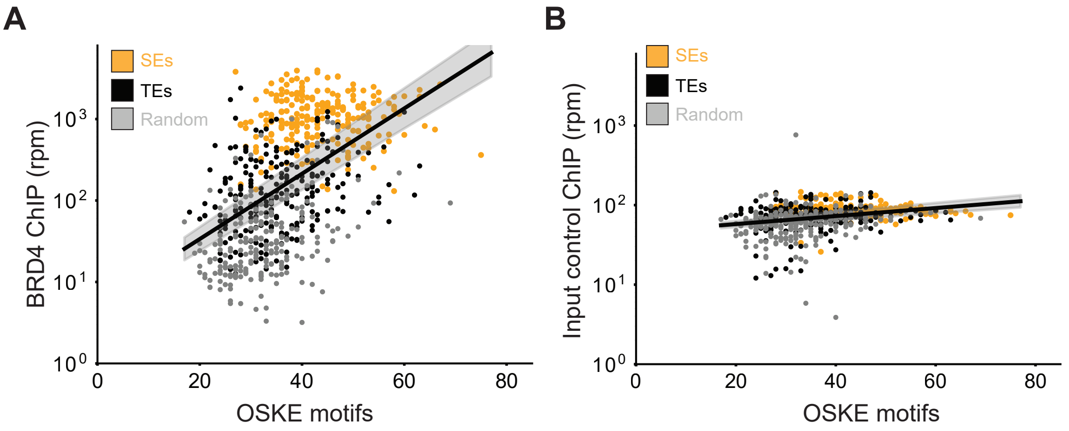


**Fig S5. Mammalian genomes show correlation between high occupancy of coactivator and motif density at regulatory elements, Related to Figure 6**

**A.** BRD4 ChIP-Seq counts (y-axis, reads-per-million) against total motifs of OCT4+SOX2+KLF4+ESRRB over 20kb regions centered on SEs (orange), TEs (black), and random loci (gray).

**B.** Same as **(A)** with data from sequenced input.

The black line represents a linear fit inferred between the logarithmic ChIP signal and motif count, and the grey shaded regions represent the 95% confidence intervals in the inferred parameters. The linear model explains a sizable fraction of the observed variance ($R^2 \approx 0.28$) for the BRD4 signal, but not for input control ($R^2 \approx 0.07$).

#### Figure S6

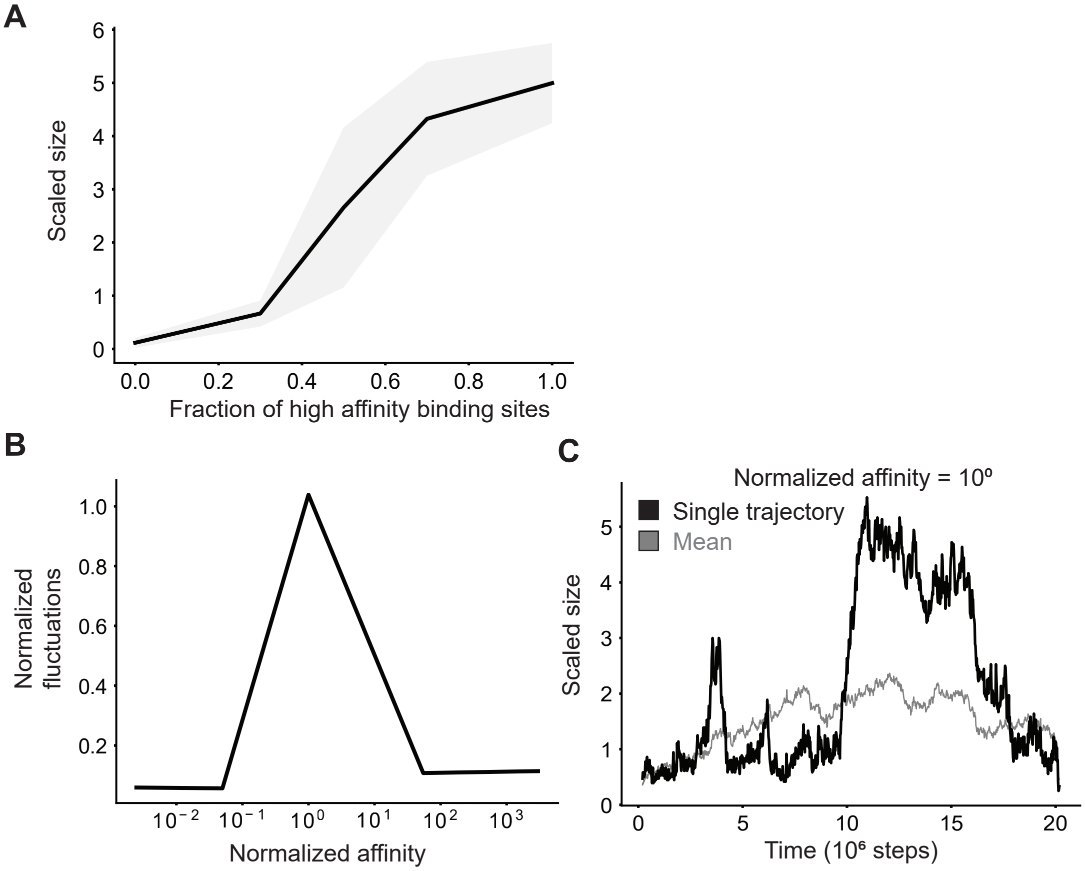


**Fig S6. Condensates form above a threshold fraction of high affinity TF-DNA sites, exhibit large fluctuations and form transient clusters at TF-DNA affinities close to the condensation threshold, Related to Figures 3,4**

**A.** Simulations predict a shift in scaled size from stoichiometric binding (≈1) to phase separation (>1) with increasing fraction of high-affinity TF binding sites on DNA. DNA is made of two types of TF binding sites, and increasing fraction along x-axis represents increase in relative proportion of high affinity binding sites.

**B.** Normalized fluctuations in scaled size (variance over mean) shows a peak with increasing normalized affinity of each binding site on DNA; affinity normalized to threshold affinity of E=12kT.

**C.** Typical simulation trajectories show dynamic formation and disassembly of clusters (Transition between low and high scaled size) at intermediate affinities.

#### SUPPLEMENTARY TABLES

| Figure label | Varied/key parameter |
|---|---|
| Fig 1B | $L = 44$ units |
| Fig S2 | $+/-$ DNA, $L = 40$ |
| Fig 2A, Fig S3C | $L = 48{,}44$ units |
| Fig 2C | $L = 28$ units |
| Fig 3A, Fig S6 | $E_{TF-DNA} = 6,9,12,16,20$ kT |
| Fig 3E | $E_{IDR-IDR} = 0,0.5,0.75,1,1.25,1.5$ |
| Fig 4A | $N_{binding-sites} = 0,10,20,30,40$ |
| Fig 3B ,Fig 3F, Fig 4B | $N_{coA} = 50,100,175,250,300,400$ |
| Fig 4F | $DNA = (D_{40}A_{20}D_{40}, (AD_4)_{20})$ |
| Fig 5A-5B | $DNA = (D_{30}A_{10}D_{20}A_{10}D_{30}, (AD_4)_{20})$ |
| Fig S6 | $N_{master} = 0,9,15,21,30$ |

**Table S1. Key simulation parameters used to generate figures in manuscript, Related to Figures 1,2,3,4,5**

Above table highlights variables changed for different simulation plots. Additional details on simulation parameters are split as:

*Explicit-IDR simulations* in Fig 1B, Fig 2A,Fig S2,S3C: Key constant parameters are $DNA = A_{10}, TF = BD_4, coactivator = CE_9$, $N_{DNA} = 1$, $N_{TF} = 100$, $N_{coA} = 200$, $E_{TF-DNA} = 20kT$, $E_{D-D} = E_{D-E} = 1\ kT, E_{D-E} = 1.25\ kT$.

*Implicit-IDR simulations* in Fig 2C, 3A-B, 3E-F, Fig 4A-B Fig S3A-B, Fig S6A-C: Key constant parameters are $DNA = A_{10}, TF = B, coactivator = C$, $N_{DNA} = 1$, $N_{TF} = 100$, $N_{coA} = 300$, $E_{TF-DNA} = 16kT$, $E_{TF-TF} = 1\ kT$, $E_{TF-coA}, E_{coA-coA} = 1.5\ kT$, $L_{box} = 28\ units$.

*Implicit-IDR large DNA simulations* in Fig 4F,5A-B: Key constant parameters are $DNA = D_{40}A_{20}D_{40}, TF = B, coactivator = C$, $N_{DNA} = 1$, $N_{TF} = 1000$, $N_{coA} = 3000$, $E_{TF-DNA} = 16kT$, $E_{TF-TF} = 1\ kT$, $E_{TF-coA}, E_{coA-coA} = 1.5\ kT$, $L_{box} = 72\ units$

| Name | Sequence |
|---|---|
| ODNA_20 | TGTAAAACGACGGCCAGTGGATCCTAGGCTTA**ATTTGCAT**TGCAGTAC**ATTTGCAT**GCATGAAT**ATTTGCAT**TAAGCTTG**ATTTGCAT**GTTTCAGA**ATTTGCAT**CGGCTAGC**ATTTGCAT**GGGCTAGA**ATTTGCAT**GCCGGATA**ATTTGCAT**GGCGATTC**ATTTGCAT**GCCAAATC**ATTTGCAT**GCATGAAC**ATTTGCAT**GGCTTACA**ATTTGCAT**GAAACATA**ATTTGCAT**CGATCGAA**ATTTGCAT**GTAGCCGA**ATTTGCAT**GTAGCTAA**ATTTGCAT**GAAATCGG**ATTTGCAT**GTAGCAAT**ATTTGCAT**CTAGCCTA**ATTTGCAT**ACCCTAGC**ATTTGCAT**TAGATTCGGCGGCCGCGTCATAGCTGTTTCCTG |
| ODNA_20sc | TGTAAAACGACGGCCAGTGGATCCTAGGCTTAATTGCCTCATCCCCTGAAATCGTTAGTGATCAGACCATTCTCTATTAATTTTAGGTGACTCTGAATCTAAATAAACATCTTTGAGATATGCTTACGATATAATGATCACTTAAGTCATCATTTGTTATCTTACAGATTTGAGATGCCAACTTTGTGGTGGCCTTAAATTGTAAGCTGAAAACCGTGAAGGAAGAGCGTTTTTGGCATATAGGTGAACTCGGTTCGTTAGCATCAGTCCGGTTCATCTGCTAGGCTGTTATCTATTATTTTATTATTCTAAATTGTGACGACGTGATAGTGGCAATCACTGACTAGATTCGGCGGCCGCGTCATAGCTGTTTCCTG |
| ODNA_5_uniform | TGTAAAACGACGGCCAGTGGATCCTAGGCTTA**ATTTGCAT**TGCAGTACATGACTCAGCATGAATAGAGTACGTAAGCTTGGTGATCACGTTTCAGA**ATTTGCAT**CGGCTAGCAGAGTACGGGGCTAGAGACTGCTAGCCGGATAGACTGCTAGGCGATTC**ATTTGCAT**GCCAAATCATGACTCAGCATGAACATGACTCAGGCTTACAGTGATCACGAAACATA**ATTTGCAT**CGATCGAAAGAGTACGGTAGCCGAGTGATCACGTAGCTAAGACTGCTAGAAATCGG**ATTTGCAT**GTAGCAATATGACTCACTAGCCTAAGAGTACGACCCTAGCGTGATCACTAGATTCGGCGGCCGCGTCATAGCTGTTTCCTG |
| ODNA_5_middle | TGTAAAACGACGGCCAGTGGATCCTAGGCTTAATCTTTAATGCAGTACATGACTCAGCATGAATAGAGTACGTAAGCTTGGTGATCACGTTTCAGATCGAAATTCGGCTAGCAGAGTACGGGGCTAGAGACTGCTAGCCGGATA**ATTTGCAT**GGCGATTC**ATTTGCAT**GCCAAATC**ATTTGCAT**GCATGAAC**ATTTGCAT**GGCTTACA**ATTTGCAT**GAAACATACCCAGTAGCGATCGAAAGAGTACGGTAGCCGAGTGATCACGTAGCTAAGACTGCTAGAAATCGGGGGTCATCGTAGCAATATGACTCACTAGCCTAAGAGTACGACCCTAGCGTGATCACTAGATTCGGCGGCCGCGTCATAGCTGTTTCCTG |

**Table S2. Annotated sequence of DNAs used in droplet assays, Related to Figures 1,2,3,4**
Sequence for each DNA species used in droplet assays. M13 (-21) and M13 reverse primer sequences used in PCR to fluorescently label and amplify DNA are underlined. The octamer motif sequence (ATTTGCAT) is bolded.

| # of binding sites | Sequence |
|---|---|
| 0 | CAGTGGATCCTAGGCTTAATTGCCTCATCCCCTGAAATCGTTA<br>GTGATCAGACCATTCTCTATTAATTTTAGGTGACTCTGAATCT<br>AAATAAACATCTTTGAGATATGCTTACGATATAATGATCACTT<br>AAGTCATCATTTGTTATCTTACAGATTTGAGATGCCAACTTTG<br>TGGTGGCCTTAAATTGTAAGCTGAAAACCGTGAAGGAAGAGC<br>GTTTTTGGCATATAGGTGAACTCGGTTCGTTAGCATCAGTCCG<br>GTTCATCTGCTAGGCTGTTATCTATTATTTTATTATTCTAAATT<br>GTGACGACGTGATAGTGGCAATCACTGACTAGATTCGGCGGC<br>CGCGTCA |
| 1 | CAGTGGATCCTAGGCTTA**ATTTGCAT**ATCCCCTGAAATCGTTA<br>GTGATCAGACCATTCTCTATTAATTTTAGGTGACTCTGAATCT<br>AAATAAACATCTTTGAGATATGCTTACGATATAATGATCACTT<br>AAGTCATCATTTGTTATCTTACAGATTTGAGATGCCAACTTTG<br>TGGTGGCCTTAAATTGTAAGCTGAAAACCGTGAAGGAAGAGC<br>GTTTTTGGCATATAGGTGAACTCGGTTCGTTAGCATCAGTCCG<br>GTTCATCTGCTAGGCTGTTATCTATTATTTTATTATTCTAAATT<br>GTGACGACGTGATAGTGGCAATCACTGACTAGATTCGGCGGC<br>CGCGTCA |
| 2 | CAGTGGATCCTAGGCTTA**ATTTGCAT**TGCAGTACATGAATCA<br>TCATGAAT**ATTTGCAT**TCTATTAATTTTAGGTGACTCTGAATC<br>TAAATAAACATCTTTGAGATATGCTTACGATATAATGATCACT<br>TAAGTCATCATTTGTTATCTTACAGATTTGAGATGCCAACTTT<br>GTGGTGGCCTTAAATTGTAAGCTGAAAACCGTGAAGGAAGAG<br>CGTTTTTGGCATATAGGTGAACTCGGTTCGTTAGCATCAGTCC<br>GGTTCATCTGCTAGGCTGTTATCTATTATTTTATTATTCTAAAT<br>TGTGACGACGTGATAGTGGCAATCACTGACTAGATTCGGCGG<br>CCGCGTCA |
| 3 | CAGTGGATCCTAGGCTTA**ATTTGCAT**TGCAGTACATGAATCA<br>TCATGAAT**ATTTGCAT**TAAGCTTGGTGATCACGTTTCAGA**ATT**<br>**TGCAT**AAACATCTTTGAGATATGCTTACGATATAATGATCACT<br>TAAGTCATCATTTGTTATCTTACAGATTTGAGATGCCAACTTT<br>GTGGTGGCCTTAAATTGTAAGCTGAAAACCGTGAAGGAAGAG<br>CGTTTTTGGCATATAGGTGAACTCGGTTCGTTAGCATCAGTCC<br>GGTTCATCTGCTAGGCTGTTATCTATTATTTTATTATTCTAAAT<br>TGTGACGACGTGATAGTGGCAATCACTGACTAGATTCGGCGG<br>CCGCGTCA |
| 4 | CAGTGGATCCTAGGCTTA**ATTTGCAT**TGCAGTACATGAATCA<br>TCATGAAT**ATTTGCAT**TAAGCTTGGTGATCACGTTTCAGA**ATT**<br>**TGCAT**CAGCTAGCAGAGTACGGGGCTAGA**ATTTGCAT**ATCAC<br>TTAAGTCATCATTTGTTATCTTACAGATTTGAGATGCCAACTTT<br>GTGGTGGCCTTAAATTGTAAGCTGAAAACCGTGAAGGAAGAG<br>CGTTTTTGGCATATAGGTGAACTCGGTTCGTTAGCATCAGTCC<br>GGTTCATCTGCTAGGCTGTTATCTATTATTTTATTATTCTAAAT<br>TGTGACGACGTGATAGTGGCAATCACTGACTAGATTCGGCGG<br>CCGCGTCA |
| 5 | CAGTGGATCCTAGGCTTA**ATTTGCAT**TGCAGTACATGAATCA<br>TCATGAAT**ATTTGCAT**TAAGCTTGGTGATCACGTTTCAGA**ATT**<br>**TGCAT**CAGCTAGCAGAGTACGGGGCTAGA**ATTTGCAT**TCCGG<br>ATAGACTGCTAGGCGATTC**ATTTGCAT**TTTGAGATGCCAACTT |

| | |
|---|---|
| | TGTGGTGGCCTTAAATTGTAAGCTGAAAACCGTGAAGGAAGAGCGTTTTTGGCATATAGGTGAACTCGGTTCGTTAGCATCAGTCCGGTTCATCTGCTAGGCTGTTATCTATTATTTTATTATTCTAAATTGTGACGACGTGATAGTGGCAATCACTGACTAGATTCGGCGGCCGCGTCA |
| 6 | CAGTGGATCCTAGGCTTA**ATTTGCAT**TGCAGTACATGAATCATCATGAAT**ATTTGCAT**TAAGCTTGGTGATCACGTTTCAGA**ATTTGCAT**CAGCTAGCAGAGTACGGGGCTAGA**ATTTGCAT**TCCGGATAGACTGCTAGGCGATTC**ATTTGCAT**GCCAAATCATGACTCAGCATGAAC**ATTTGCAT**TGTAAGCTGAAAACCGTGAAGGAAGAGCGTTTTTGGCATATAGGTGAACTCGGTTCGTTAGCATCAGTCCGGTTCATCTGCTAGGCTGTTATCTATTATTTTATTATTCTAAATTGTGACGACGTGATAGTGGCAATCACTGACTAGATTCGGCGGCCGCGTCA |
| 7 | CAGTGGATCCTAGGCTTA**ATTTGCAT**TGCAGTACATGAATCATCATGAAT**ATTTGCAT**TAAGCTTGGTGATCACGTTTCAGA**ATTTGCAT**CAGCTAGCAGAGTACGGGGCTAGA**ATTTGCAT**TCCGGATAGACTGCTAGGCGATTC**ATTTGCAT**GCCAAATCATGACTCAGCATGAAC**ATTTGCAT**GGCTTACAGTGATCACGAAACATA**ATTTGCAT**TTGGCATATAGGTGAACTCGGTTCGTTAGCATCAGTCCGGTTCATCTGCTAGGCTGTTATCTATTATTTTATTATTCTAAATTGTGACGACGTGATAGTGGCAATCACTGACTAGATTCGGCGGCCGCGTCA |
| 8 | CAGTGGATCCTAGGCTTA**ATTTGCAT**TGCAGTACATGAATCATCATGAAT**ATTTGCAT**TAAGCTTGGTGATCACGTTTCAGA**ATTTGCAT**CAGCTAGCAGAGTACGGGGCTAGA**ATTTGCAT**TCCGGATAGACTGCTAGGCGATTC**ATTTGCAT**GCCAAATCATGACTCAGCATGAAC**ATTTGCAT**GGCTTACAGTGATCACGAAACATA**ATTTGCAT**CGATCGAAAGAGTACGGTAGCCGA**ATTTGCAT**CAGTCCGGTTCATCTGCTAGGCTGTTATCTATTATTTTATTATTCTAAATTGTGACGACGTGATAGTGGCAATCACTGACTAGATTCGGCGGCCGCGTCA |
| 5 (with lower density) | CAGTGGATCCTAGGCTTA**ATTTGCAT**TGCAGTACATGACTCAGCATGAATAGAGTACGTAAGCTTGGTGATCACGTTTCAGA**ATTTGCAT**CGGCTAGCAGAGTACGGGGCTAGAGACTGCTAGCCGGATAGACTGCTAGGCGATTC**ATTTGCAT**GCCAAATCATGACTCAGCATGAACATGACTCAGGCTTACAGTGATCACGAAACATA**ATTTGCAT**CGATCGAAAGAGTACGGTAGCCGAGTGATCACGTAGCTAAGACTGCTAGAAATCGG**ATTTGCAT**GTAGCAATATGACTCACTAGCCTAAGAGTACGACCCTAGCGTGATCACTAGATTCGGCGGCCGCGTCA |

**Table S3. Annotated sequence of DNAs used in luciferase reporter assays, Related to Figure 4**

Sequences tested in luciferase reporter assays are provided here with the octamer motif sequence (ATTTGCAT) bolded. The sequences provided were cloned into the SalI site of the previously characterized pGL3-basic OCT4 (Whyte et al 2013) as described in methods.

#### Supplementary Movie Captions

**Movie S1. Simulation trajectory of phase separation mediated by DNA at low protein concentrations, Related to Figure 2**

Typical simulation trajectory of DNA-scaffolded condensate formation and subsequent disassembly (sub-titles) at low protein concentrations (Ref Fig 2A). DNA particles are in red, TFs in yellow, coactivators in blue (same as Fig 1A), and all IDRs are in grey. The movie is played at a 10x acceleration in the frame-rate.

**Movie S2. Simulation trajectory of phase separation mediated by DNA at high protein concentrations, Related to Figure 2**

Typical simulation trajectory of DNA-scaffolded condensate formation and ejection of DNA after disruption of TF-DNA interactions(sub-titles) at high protein concentrations (Ref Fig 2A). DNA particles are in red, TFs in yellow, coactivators in blue (same as Fig 1A), and all IDRs are in grey. The movie is played at a 10x acceleration in the frame-rate.
